# Cytomegalovirus promotes proliferation and survival of prostate cancer cells and constitutes a therapeutic target

**DOI:** 10.1101/2023.10.01.560348

**Authors:** Johanna Classon, Moa Stenudd, Margherita Zamboni, Kanar Alkass, Carl-Johan Eriksson, Lars Pedersen, Alrik Schörling, Anna Thoss, Anders Bergh, Pernilla Wikström, Hans-Olov Adami, Henrik T Sørensen, Henrik Druid, Jonas Frisén

## Abstract

Metastatic prostate cancer is incurable and new therapeutic targets and drugs are needed. Viruses are associated with several cancer types, but their connection to prostate cancer is unclear. Here we show that human herpes virus cytomegalovirus (CMV) infection is common in the healthy prostate epithelium as well as in prostate cancer, with 85% of tumors being infected to varying degrees. The CMV gene locus *UL122-UL123* upheld viral genome persistence in endogenously CMV infected prostate cancer cell lines. CMV promoted prostate cancer cell viability independently of androgen receptor status and anti-androgen resistance, partly through CMV *UL97* and the androgen signaling pathway. DNA intercalation mitigated CMV infection and reduced CMV-dependent tumor size in xenotransplantation experiments. The anti-herpes drug aciclovir showed modest effects, but the well tolerated CMV UL97 kinase inhibitor maribavir partly mimicked CMV loss by inducing apoptosis and attenuating proliferation, resulting in reduced tumor growth *in vivo*. We conclude that CMV infects prostate cells *in vivo* and alters core prostate cancer cell properties, suggesting that it can be therapeutically targeted to improve prostate cancer outcomes.

---

Prostate cancer is one of the most common tumor types with more than 350 000 prostate cancer deaths reported worldwide each year, making prostate cancer one of the leading causes of death from cancer. Standard of care for advanced prostate cancer patients is androgen deprivation therapy and chemotherapy. Although new therapies that prolong life have emerged the last few years, including poly (ADP-ribose) polymerase and prostate-specific membrane antigen targeted therapies, metastatic prostate cancer is still incurable^1^ and it is important to identify new therapeutic targets.

A person is estimated to carry multiple chronic viral infections^2^, which may have profound effects on the host, as they have the potential to modify cellular functions and the immune system^3^. Chronic viral infections can induce cancer, alter disease progression and may themselves be therapeutic targets. For example, human papillomavirus causes cervical cancer and can be targeted by vaccination to decrease cancer incidence^4,5^.

Human cytomegalovirus (CMV) is a herpes virus, which the majority of humans are chronically infected by, indicated by 83% having antibodies against the virus^6^. This is likely a substantial underestimation of the proportion of the population that is infected, as it is very common to detect CMV DNA in seronegative individuals^7–14^, indicating that they are infected but lack a detectable antibody response. CMV seropositivity is associated with increased general cancer mortality in men^15^, but not associated with prostate cancer incidence^16,17^. We reported that local T cell immunity to CMV in HLA-A*02:01 prostate cancer patients is associated with disease recurrence after prostatectomy^18^. Moreover, we recently found that CMV seropositivity is associated with increased prostate cancer mortality^17^. These studies suggest that, although CMV does not appear to cause prostate cancer^16,17^, either CMV or an immune response to CMV is associated with a more aggressive disease. Here we explore CMV infection characteristics in the prostate, the role of CMV in models of prostate cancer and its potential as a therapeutic target in advanced prostate cancer.

We report that CMV infection is common in epithelial cells of the healthy prostate as well as in prostate cancer. We find that commonly studied prostate cancer cell lines are CMV infected, and that CMV promotes cell survival and proliferation, identifying CMV as a therapeutic target in castration-sensitive and -resistant prostate cancer. We demonstrate that several pharmaceutical compounds targeting CMV, which are either used in clinical practice or recently approved, have anti-cancer effects in prostate cancer models.

## RESULTS

### CMV in epithelial cells in the healthy prostate

It is well established that CMV persists in hematopoietic lineage cells^19,20,8^, from which reactivation events are thought to occur throughout life^21^. Several organs containing specialized epithelial cells have also been suggested as chronic CMV infection sites^12,22,23^, including the normal and cancerous prostate gland^24–26^. It is, however, technically challenging to detect chronic/latent infection, which has led to conflicting data and hampered research on CMV in tissues in health and diseases^27^.

We examined to what extent CMV is present in the human prostate with several orthogonal methods. The 235 kbp long CMV genome consists of more than 150 genes. Compared to in active/lytic infection, viral genes in chronic/latent CMV infection are as broadly expressed but at much lower levels^23,28^. We first examined CMV protein expression, with the presumption that these may be more accessible for detection than other viral products. We detected the well-characterized CMV proteins UL97 and US28 in prostate tissue homogenates by immunoblot (n=8; Fig. 1A).

**Figure 1:**
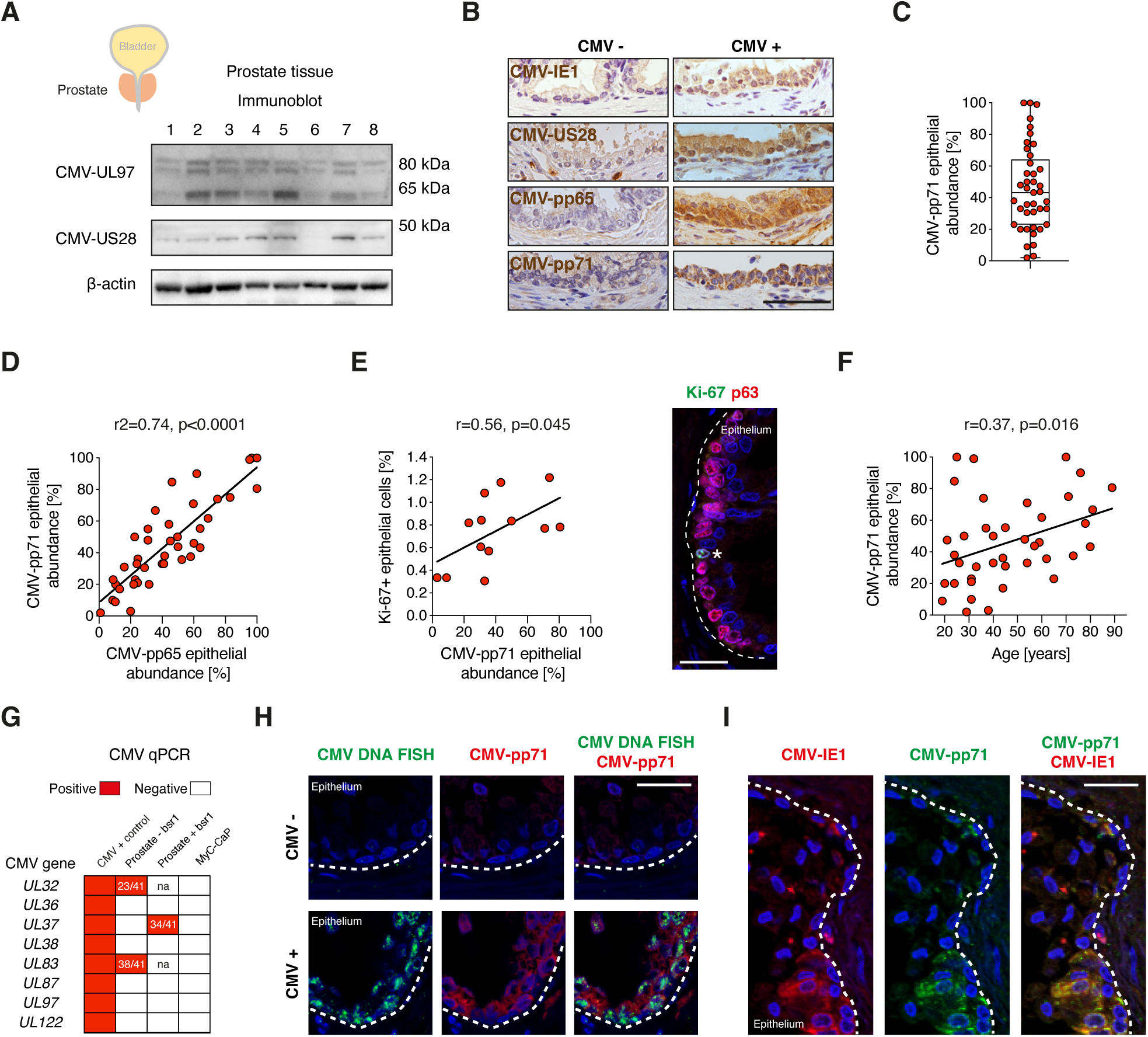
CMV in prostate epithelial cells. **A)** Immunoblot of prostate tissue homogenates for the proteins CMV-UL97 (with three bands as expected^71^), CMV-US28 and β-actin from eight men. kDa is kilodalton. **B)** Images of prostate sections with CMV antibody staining in brown in the epithelium. Nuclei are labeled in purple with hematoxylin. Scale bar 50 μm. **C)** Quantification of CMV-pp71 abundance in epithelial cells. All datapoints (n=41) are shown in graph and summarized in box plot (median, 25-75^th^ percentiles, error bars show min-max values). **D)** Linear regression comparing CMV-pp71 and CMV-pp65 abundance in the prostate epithelium (n=41). **E)** Pearson correlation of percentage of Ki-67^+^ cells and CMV-pp71 abundance (n=13). Labeling with Ki-67 (green) and p63 (red). Star points out a Ki-67^+/^p63^+^ epithelial cell. Scale bar: 25 μm. **F)** Pearson correlation between CMV-pp71 epithelial abundance and age (n=41). **G)** CMV qPCR heatmap. CMV qPCR assays detected CMV DNA in positive control (purified CMV DNA) but not in negative control (mouse prostate cancer cell line MyC-CaP). Numbers denote how many prostates were positive and evaluated with qPCR with or without *Bsr1* restriction enzyme pre-treatment. Three prostate samples were evaluated in screening of qPCR assays. Na is not analyzed. **H)** Co-labeling of CMV DNA (green) and CMV-pp71 (red). Scale bar 25 μm. **I)** Co-labeling of CMV-pp71 (green) and CMV-IE1 (red). Scale bar 25 μm. In (E), (H) and (I), nuclei are labeled in blue with DAPI and dotted lines depict the basal lamina of the epithelium.

CMV has broad cellular tropism. To localize CMV within prostates, we turned to immunohistochemistry (IHC), which allowed us to characterize and quantify CMV protein expression among cell types. We focused on characterizing CMV infection in epithelial cells since these are the cells of origin of prostate cancer. The four CMV proteins IE1, US28, pp65 and pp71 were detected in epithelial areas (Fig. 1B-C) with high concordance (Fig. 1D, Figure S1A-B). No signal was detected in negative control assays (Fig. S1C) and all CMV IHC assays identified CMV in positive controls (Fig. S1D). CMV was detected in the prostate in all subjects, with the infected proportion of the prostate epithelium ranging from 2% to 100% with a mean of 46% (n=41, Fig. 1C). Within CMV^+^ epithelial areas, either patches of cells or whole areas of glandular structures were infected (Fig. 1B), that involved one or more of the main epithelial subtypes basal-, luminal-and neuroendocrine cells (Fig. S1E). Interestingly, CMV abundance correlated with the number of epithelial cells that were positive for the proliferation marker Ki-67 (r=0.56, p=0.045, Fig. 1E) and the proportion of CMV infected epithelial cells increased with age (r=0.37, p=0.016, Fig. 1F), possibly reflecting expansion of CMV infected cells.

CMV DNA and RNA are easily found during active/lytic infection but more challenging to detect when the virus is chronic/latent. In prostates, expression of CMV genes has not been detected by RNA-sequencing^23,29,30^, confirmed in an analysis of deeply sequenced prostate cancer (see Online Methods). This finding implies a chronic/latent nature of CMV infection in the prostate. We next analyzed presence of CMV DNA. As a positive control, we detected CMV DNA with two qPCR assays (CMV-*UL32*, CMV-*UL83*) in CD14 enriched peripheral blood mononuclear cells (latent infection; Fig. S2A-D) independent of CMV serostatus but, as expected^31–33^, CMV IgG^+^ donors had highest quantities of CMV (Fig. S2D). With these qPCR assays, CMV was also identified in the prostate, and the only CMV qPCR^-^ prostate in this cohort had the lowest CMV epithelial abundance (2%) by IHC (Fig. 1C, 1G, Fig. S1A). Sequencing of PCR products in positive assays confirmed that they represented the correct CMV genes (Fig. S2E). Speaking for a latent, difficult to detect, CMV infection, six other CMV qPCR assays, that readily identified purified CMV DNA, did not identify CMV in prostates, but one (CMV-*UL37*) revealed CMV after pre-treatment of DNA with a restriction enzyme (Fig. 1G). Difficulties in capturing latent CMV by standard PCR assays was expected^20,34^, and may be related to complex viral DNA conformations. CMV DNA (*RNA2.7* gene) was furthermore detected in tissue sections by in situ hybridization and was localized exclusively in CMV protein positive cells (Fig. 1H), validating presence of CMV nuclei acids and viral gene products in prostate epithelial cells.

Several lines of evidence support the specificity of our CMV detection in the prostate. First, we detected several different CMV proteins, and their abundance correlated (Fig. 1D, Fig. S1B). Second, different CMV proteins were co-detected in the same prostate epithelial cells (Fig. 1I, Fig. S1F). Third, CMV DNA could be amplified from prostate tissue (Fig. 1G). Fourth, CMV DNA detected by in situ hybridization, was selectively present in CMV protein positive cells (Fig. 1H). Taken together, these findings demonstrate that CMV infection is common and widespread in the prostate.

### CMV in prostate cancer

CMV seropositivity does not increase the risk of developing prostate cancer^16,17^, but is associated with prostate cancer mortality^17^, suggesting that CMV may have direct effects if present in tumor cells. We analyzed the presence and abundance of CMV in 20 tumors in prostatectomy specimens from patients with prostate cancer (Fig. S3A).

CMV was detected by CMV pp71 IHC in 17/20 (85%) tumors, with ten having very high abundance of CMV (90-100% of cancer cells infected, Fig. 2A). In prostate samples from three patients with CMV negative tumors, CMV was also absent from the benign epithelium (Fig. S3B). We also found incidental prostate cancer in two post-mortem donors, of which one was CMV^+^ and the other was CMV^-^, as determined with CMV DNA in situ hybridization and IHC (Fig. S3C). These results show that prostate cancer cells often, but not always, are infected by CMV, supporting that CMV may not be essential in the etiology of prostate cancer.

**Figure 2:**
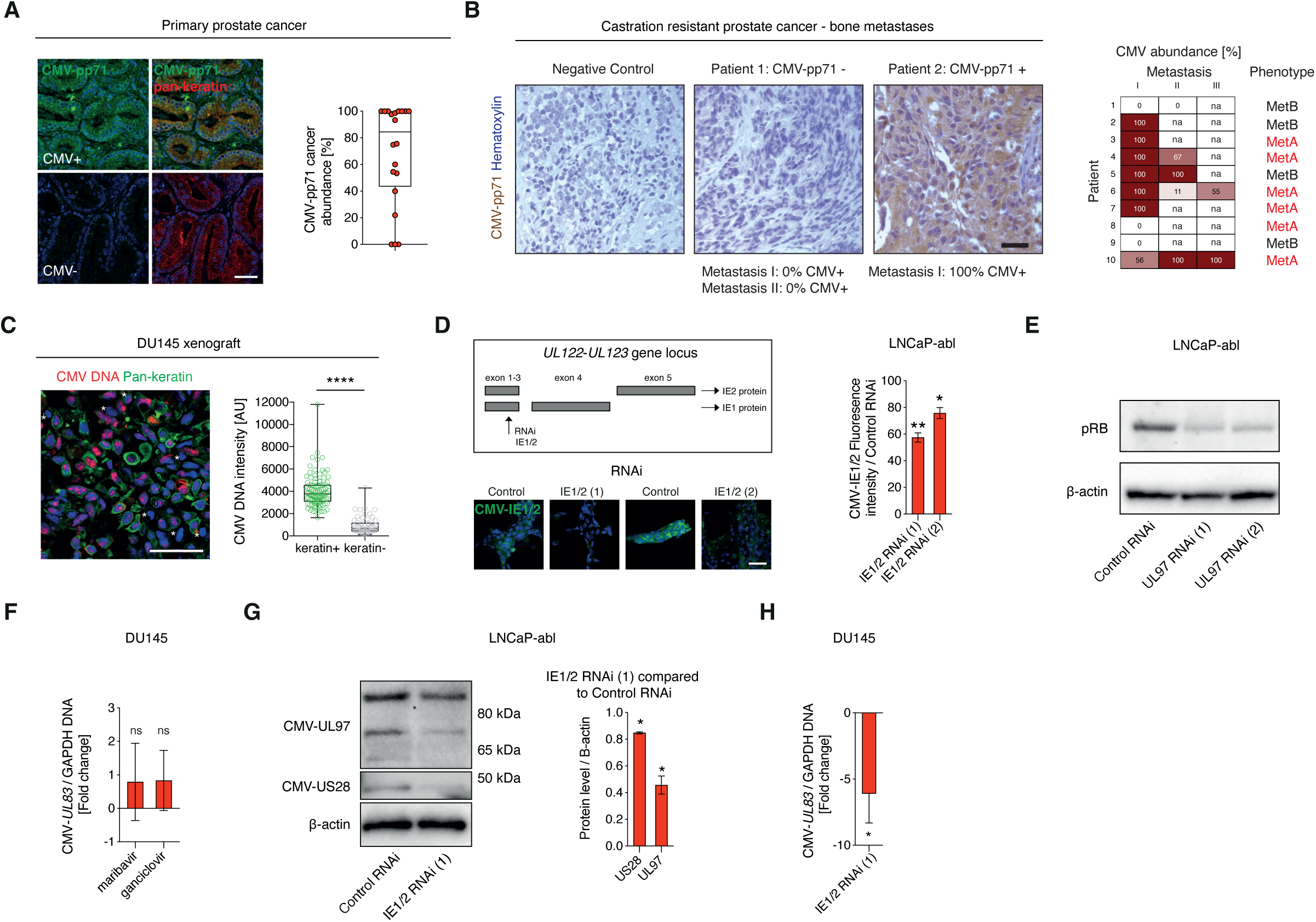
CMV in prostate cancer is maintained by the CMV locus *UL122-UL123*. **A)** Abundance of CMV-pp71 (%) in tumors (n=20), determined by IHC. Pan-keratin was used as a marker for benign and malignant epithelial cells. Prostate cancer was identified histologically in pathologist-outlined areas. Scale bar: 25 μm. **B)** Bone metastases from 10 CRPC patients were examined for CMV-pp71 protein expression with IHC. Nuclei are labeled in purple with hematoxylin. Scale bar: 50 μm. Phenotypes were defined as MetA (high AR activity), MetB (dedifferentiated phenotype) or MetC (EMT phenotype)^35^. In this cohort, metastases with MetA and MetB phenotypes were present. In five patients, more than one metastasis was examined, labelled metastases I, II and III and percentage of CMV-pp71+ areas are shown in heat map. Na is not analysed. **C)** CMV DNA FISH in DU145 xenografts (n=3). Fluorescence intensity in cell nuclei in pan-keratin^+^ and pan-keratin^-^ cells were compared with un-paired t-test. AU = arbitrary unit. Scale bar: 50 μm. Pan-keratin was used as a marker of epithelial cells. All datapoints (all cells analyzed) are shown in plot. **D)** LNCaP-abl transfected with IE1/2 RNAi (1) or IE1/2 RNAi (2), targeting the *UL122-UL123* exon 3 (illustration), reduced mean IE1/2 fluorescence compared to Control RNAi, analyzed with one-sample t-test. Scale bar: 50 μm **E)** pRB and ý-actin expression was examined by immunoblot. **F)** DU145 treated with vehicle, 30 μM maribavir or 100 μM ganciclovir examined with CMV-*UL83* qPCR after three days (n=4). Data is shown as fold change and examined with paired t-tests. **G)** Immunoblot of LNCaP-abl four days after transfection with control RNAi, IE1/2 RNAi (1). CMV-US28, CMV-UL97 and ý-actin expression was examined, quantified (n=3) and analyzed with one-sample t-test. kDa is kilodalton. **H)** CMV DNA abundance determined with CMV-*UL83* qPCR in DU145 four days after transfection with IE1/2 RNAi (1) compared to control RNAi (n=6). Data is displayed as negative fold change. Box plots in (A) and (C) show median; boxes 25-75 percentiles and error bars are max and min. p<0.05 is*, p<0.01 is **, p<0.0001 is ****. Ns is non-significant. Nuclei are labeled in blue with DAPI in (A), (C) and (D).

Primary prostate cancer cells often metastasize to distal sites such as bone. We evaluated metastases from ten patients in a separate cohort with CMV-pp71 IHC to examine the distribution of CMV. CMV was detected in metastases in 70% of patients, with a total of 9/13 (69%) CMV^+^ metastases examined being homogenously infected (Fig. 2B, Fig. S4A). Four examined bone metastases from patients with castration resistant prostate cancer (CRPC) were positive for CMV-*UL37* qPCR. CMV was not limited to a particular metastasis phenotype but could be present in metastases with high androgen receptor (AR) activity (MetA) or a dedifferentiated phenotype (MetB)^35^ (Fig. 2B). The observation that CMV is commonly present in primary prostate cancer and prostate cancer metastases, together with an increased prostate cancer mortality among CMV seropositive prostate cancer patients^17^, suggest that CMV may exert direct cancer promoting functions.

### CMV in prostate cancer cell lines

Direct functions of CMV on prostate cancer biology can be studied in experimental systems. Infecting prostate cancer cells with purified CMV virions may not accurately mimic latent in vivo prostate cancer infection. We therefore assessed whether cell lines derived from patients with advanced prostate cancer are endogenously CMV infected, thereby offering a natural experimental system. Notably, all seven analyzed commonly studied prostate cancer cell lines (LNCaP, LAPC-4, VCaP, DU145, PC3, LNCaP-abl, LREX’) carried CMV, as indicated by being CMV-*UL83* qPCR^+^ (Fig. S5A). Cell lines were also CMV UL97 and US28 protein positive (Fig. S5B). CMV DNA was detected in most pan-keratin^+^ prostate cancer cells when implanted in mice as xenografts, but human CMV was, as expected, absent from mouse cells (pan-keratin^-^, Fig. 2C) and the mouse prostate cancer cell line MyC-CaP (CMV-*UL83* qPCR negative; Fig. 1G).

Even though we were unable to detect CMV RNA by RT-qPCR (Fig. S5C), gene specific RNA interference (RNAi) against the CMV genes *UL97*, *US28* and *UL122-UL123* reduced expression of the corresponding proteins, validating CMV gene expression in prostate cancer cell lines (Fig. 2D, Fig. S5D-G). Furthermore, UL97 kinase phosphorylates the Retinoblastoma protein (Rb)^36^ and *UL97* RNAi or treatment with the UL97 kinase inhibitor maribavir both resulted in reduced phosphorylation of Rb on Ser^807^/Ser^811^ in LNCaP and LREX’ (Fig. 2E, S5H), demonstrating functional activity of UL97 in prostate cancer cells. These findings show that viral protein expression and functions can be altered by specifically inhibiting them and validate that CMV is indeed present in prostate cancer cell lines.

Anti-herpes virus treatment with ganciclovir or maribavir reduce CMV abundance by blocking viral DNA synthesis and, in addition, maribavir inhibits virion assembly during virus production. Neither ganciclovir nor maribavir reduced the abundance of CMV in prostate cancer cells (Fig. 2F), suggesting that CMV is latent and consequently unaffected by these anti-viral drugs. *UL122-UL123* is an essential and well-characterized CMV gene locus encoding the proteins immediate early 1 and 2 (IE1/2), which regulate the viral life cycle^37^ and that is necessary for latent CMV genome replication^38^. *UL122-UL123* RNAi (IE1/2 RNAi) reduced expression of IE1/2 (Fig. 2D, Fig. S5G). It also reduced the expression of other CMV proteins (Fig. 2G, Fig. S5I), which we hypothesized were due to IE1/2 maintaining CMV latent genomes. Indeed, CMV DNA abundance was decreased more than five-fold in comparison to cellular DNA by IE1/2 RNAi (Fig. 2H), establishing that *UL122-UL123* is required to maintain CMV infection in prostate cancer cell lines. In summary, these findings expand upon known functions of *UL122-UL123* and suggest that latent CMV can be removed from infected cells by targeting this gene locus.

### CMV promotes prostate cancer cell survival, proliferation and androgen receptor expression

We next assessed the influence of CMV on cellular functions in prostate cancer cell lines, to determine its importance to prostate cancer biology. The CMV^+^ cell lines we analyzed represent *in vitro* models of three common phenotypes of advanced cancer: castration sensitive prostate cancer (CSPC; LNCaP, LAPC-4, VCaP), adenocarcinoma differentiated castration resistant prostate cancer (CRPC-Adeno; LNCaP-abl, LREX’) and stem cell like CRPC (CRPC-SCL; DU145, PC3). We used IE1/2 RNAi, which reduced CMV protein levels and viral load (Fig. 2C-E, 2G-H, Fig. S5D-I), to assess the influence of CMV. IE1/2 RNAi reduced viability of prostate cancer cells of all three phenotypes (Fig. 3A, Fig. S5J) and increased levels of the apoptosis marker cleaved caspase-3 in 5/7 (71%) of the examined prostate cancer cell lines, independent of castration resistant or sensitive properties (Fig. 3B). As expected, apoptosis was not induced by IE1/2 RNAi in a human CMV^-^ mouse prostate cancer cell line (Fig. 3B). IE1/2 RNAi reduced the proportion of cells being positive for the cell proliferation marker Ki-67 in CSPC and CRPC-Adeno cell lines, but not in cells of a CPRC-SCL phenotype (Fig. 3C). Since reducing CMV in prostate cancer cell lines decreases cell survival and proliferation, we conclude that CMV promotes these key prostate cancer traits.

**Figure 3:**
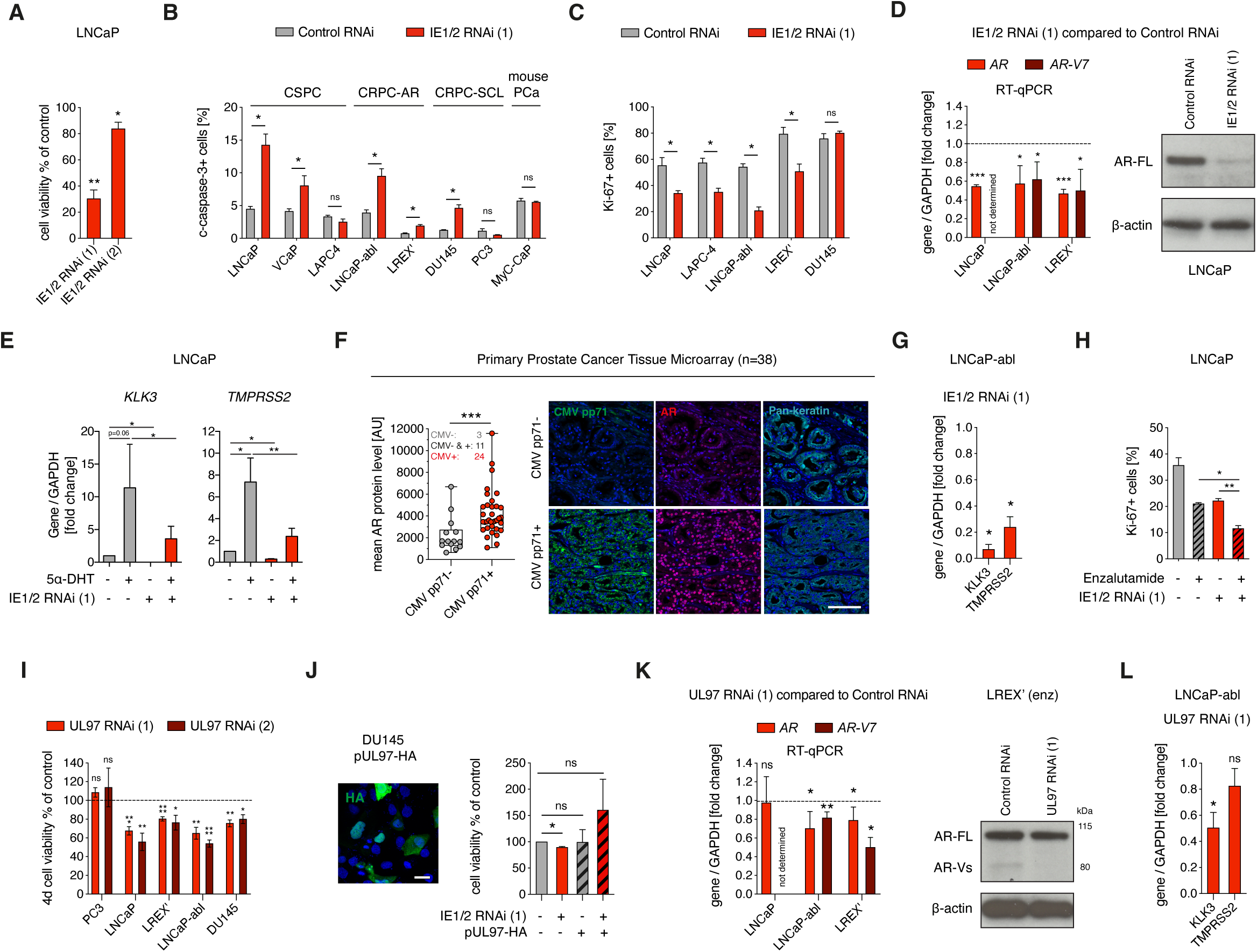
Endogenous CMV promotes cell viability and androgen receptor expression in prostate cancer. **A)** Cell viability in LNCaP four days after transfection with IE1/2 RNAi (1) (n=6) or IE1/2 RNAi (2) (n=4) compared to control RNAi. **B)** Cells transfected with control RNAi or IE1/2 RNAi (1) analyzed after three or four days for % cleaved casaspe-3+ cells (n=3-4). Phenotype of cell lines are shown in figure: CSPC, CRPC-AR and CRPC-SCL. MyC-CaP is a mouse prostate cancer (mouse PCa) cell line. **C)** Cells transfected with control RNAi or IE1/2 RNAi (1) and analyzed after three or four days for percentage of Ki-67^+^ cells (n=3). **D)** Cells transfected with control RNAi or IE1/2 RNAi (1) was examined for *AR* and *AR-V7* gene expression with RT-qPCR (n=3-4) and full length AR (AR-FL) protein expression with western blot three days after transfection. **E)** LNCaP grown in charcoal stripped FBS and transfected with control RNAi or IE1/2 RNAi (1) four days (n=3). Cells were treated with 10 nM 5α-dihydrotestosterone (DHT) one day after which gene expression of the AR target genes *TMPRSS2* and *KLK3* was examined**. F)** An association between CMV pp71 presence in cancer cells and AR protein levels in nuclei (AU; arbitrary unit) was examined in a tissue microarray with primary prostate cancer (n=38). Pan-keratin was used as a marker of epithelial cells. Each dot in graph is mean AR protein level of all CMV pp71-cells (grey) in one tumor and CMV pp71+ cells (red) in one tumor, compared with Mann-Whitney test. Three samples contained no CMV+ tumor cells, 11 samples contained both CMV+ and CMV-tumor cells and 24 samples contained only CMV+ tumor cells. Box plot shows median; box shows 25-75 percentiles and error bars show max and min. Scale bar: 100 μm. **G)** Gene expression analysis in IE1/2 RNAi (1) treated LNCaP-abl compared to control RNAi four days after transfection (n=3). **H)** Percentage of Ki-67+ cells in LNCaP transfected with control RNAi or IE1/2 RNAi (1) and treated with vehicle or 10 μM enzalutamide the following day (n=3). **I)** Cell viability evaluated four days after transfection with control RNAi, UL97 RNAi (1) or UL97 RNAi (2) (n=4-6). J) DU145 transfected with a hemagglutinin tagged UL97 plasmid (pUL97-HA). Transfection efficiency was visualized with HA detection by immunofluorescence. Scale bar is 25 μm. DU145 cells transfected with IE1/2 RNAi (1) or control RNAi was transfected with pUL97-HA or control plasmid the following day. Cell viability was evaluated after four days (n=3). K) Cells were examined for *AR* and *AR-V7* gene expression four days after transfection with Control RNAi or UL97 RNAi (1), LREX’ n=3. LNCaP n=3, LNCaP-abl n=5. AR-FL and AR-Vs protein expression four days after transfection in LREX’ was examined with western blot. kDa is kilodalton. L) *TMPRSS2* and *KLK3* expression examined after four days treatment with UL97 RNAi (1) compared to control RNAi in LNCaP. Cell nuclei in (F) and (J) are labeled in blue with DAPI. Data in bar graphs are shown as mean. Error bars in bar graphs are SEM. RT-qPCR results are shown as fold change of control. (I) and (J) were examined with one sample t-test. Unless otherwise stated, treatment effects were examined with paired t-tests. p<0.05 is*, p<0.01 is **, p<0.001 is ***, p<0.0001 is ****, ns is non-significant.

Androgen signaling is a core prostate cancer pathway that promotes growth in CSPC and CRPC-Adeno, but not in AR independent CRPC-SCL, in which AR signaling is low or absent^39^. To assess whether CMV can influence androgen signaling, we transfected CSPC cells with IE1/2 RNAi. This resulted in reduced *AR* gene expression and AR protein levels (Fig. 3D), as well as in reduced expression of the androgen regulated genes *KLK3* and *TMPRSS2* (Fig. 3E). Moreover, we found that CMV infected cells (CMV-pp71^+^) expressed higher levels of AR protein than uninfected (CMV pp71^-^) cells in primary prostate cancer samples (n=38) (Fig. 3F), suggesting that CMV increases AR levels in prostate cancer cells also *in vivo*. In LNCaP, IE1/2 RNAi reduced the number of Ki-67^+^ cells with a similar magnitude as the AR inhibitor enzalutamide and combination treatment with IE1/2 RNAi and enzalutamide further reduced proliferation compared to either treatment alone (Fig. 3G). This result implies that CMV inhibition may have additive effects to standard anti-androgen therapy.

Increased activity of AR and the AR splice variant AR-V7 by higher expression or gain of function mutations is a hallmark of CRPC-Adeno, and may act to confer resistance to anti-androgen therapy^40^. These processes are mimicked in LNCaP-abl, grown in the absence of androgens, and LREX’, grown in the presence of enzalutamide. IE1/2 RNAi decreased protein levels and expression of *AR* and *AR-V7* and their target genes *KLK3* and *TMPRSS2* in CRPC-Adeno cells (Fig. 3D, 3H), demonstrating that CMV inhibition can further attenuate androgen signaling in androgen deprived conditions.

When performing CMV-UL97 knock down experiments, we noted that several prostate cancer cell lines responded with altered cell viability. Surprisingly, we found that CMV-UL97, which does not regulate CMV latency (Fig. 2F), promotes growth in CMV dependent cell lines as knock down of CMV-UL97 reduced their cell viability (Fig. 3I) and ectopic expression of UL97 rescued CMV attenuated DU145 cells (Fig. 3J). As for IE1/2 RNAi, the effect of UL97 RNAi on cell viability was not limited to a particular prostate cancer phenotype (Fig. 3I). Furthermore, UL97 RNAi reduced *AR* and *AR-V7* expression in LNCaP-abl and LREX’, but not in LNCaP (Fig. 3K) and reduced the AR target gene *KLK3* but not *TMPRSS2* in LNCaP-abl (Fig. 3L). We conclude that UL97 may partly regulate CMV’s effects on androgen signaling and can promote prostate cancer cell viability independent of AR.

### Aciclovir in prostate cancer models and in a large population cohort

A case study reported that two prostate cancer patients with shingles (caused by the herpes virus varicella zoster) treated with valaciclovir, an aciclovir pro-drug for oral administration, for two weeks obtained a marked and lasting drop in serum prostate specific antigen^41^. As aciclovir also inhibits CMV, we wanted to assess whether the effects described in the case report could be attributed to an effect on CMV in prostate cancer. Aciclovir is a common nucleotide analogue anti-herpes virus drug, which is activated through its phosphorylation by viral proteins such as CMV UL97 kinase^42,43^. We found that CMV UL97 kinase is expressed and functioning in prostate cells (Fig. 1, 2). In addition to inhibition of viral DNA synthesis, activated aciclovir can also inhibit human DNA polymerase with low affinity, causing DNA chain termination, DNA damage and ultimately cell death^44^ (Fig. 4A). This feature of nucleotide analogues such as aciclovir can be used to target cancer cells.

**Figure 4:**
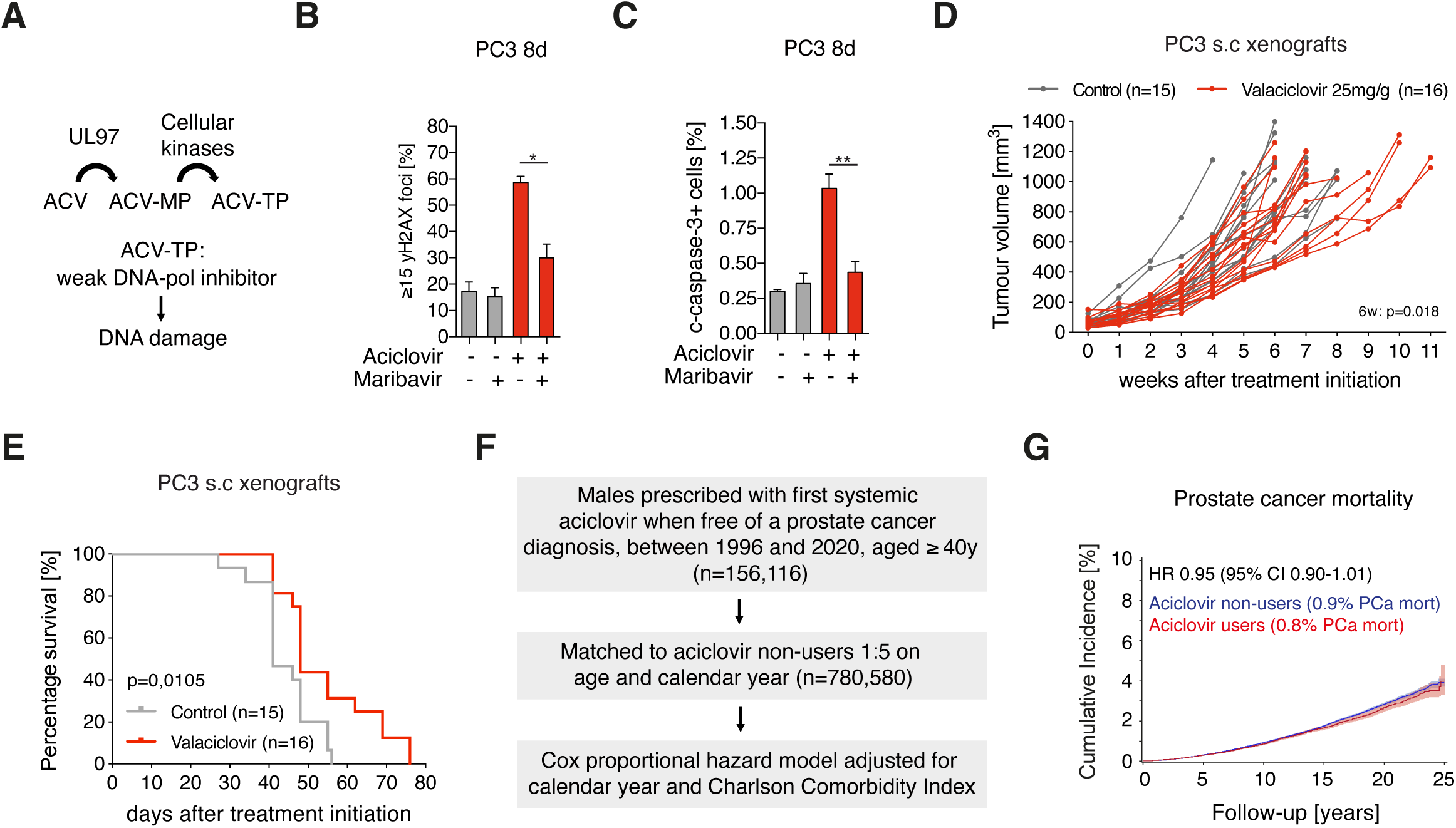
Aciclovir has modest effects against prostate cancer in models and no association with prostate cancer mortality. **A)** Illustration of aciclovir (ACV) activation and mechanism of action of activated aciclovir. ACV-MP = aciclovir monophosphate, ACV-TP = aciclovir triphosphate, DNA-pol = cellular DNA polymerase. **B-C)** In PC3, maribavir treatment (10 μM) reduced the ability of aciclovir to promote ψH2AX foci (n=3) (B) and to induce apoptosis (n=3) (C), as analyzed with paired t-tests. **D)** PC3 xenograft volume over time (weeks after treatment initiation) in control (n=15) and valaciclovir treated animals (n=16). Repeated measurement mixed-effect analysis was used to compare tumor volume over time and multiple comparison statistics was assessed at 6 weeks. **E)** Kaplan-Meier survival curve comparing survival of control (n=15) and valaciclovir (n=16) treated animals. Survival was compared using log-rank test. **F)** Flow chart of study design of aciclovir population cohort and prostate cancer mortality. **G)** Cumulative incidence (%) of prostate cancer mortality in aciclovir non-users (blue line) and aciclovir users (red line). Shaded blue and red areas represent 95% confidence intervals (CI). HR: Hazard Ratio. P<0.05 is*, p<0.01 is **.

Aciclovir (300 μM), and the related drug ganciclovir (100 μM), increased the number of γH2AX foci per nuclei (Fig. 4B, Fig. S6A-B), a marker of double stranded DNA damage and apoptosis in all seven prostate cancer cell lines examined (Fig. 4C, Fig. S6D-F). This response was abolished when CMV-UL97 was inhibited (Fig. 4B-C, S6B-F) or upon IE1/2 RNAi (Fig. S6G), demonstrating that this effect was dependent on CMV rather than being mediated by other herpes viruses or other mechanisms. Furthermore, as expected, apoptotic effects of aciclovir were dependent on active proliferation (Fig. S6H). Aciclovir induced an apoptotic response in LAPC-4 and PC3, cell lines that do not undergo apoptosis upon IE1/2 RNAi, suggesting that it does not act through direct CMV inhibition but rather induce DNA damage in cellular DNA.

Treatment with aciclovir for five days followed by seven days of no treatment enhanced its apoptosis inducing effect (Fig. S6I-J). Intermittent treatment was accompanied by an increase in cell proliferation (Fig. S6K) and DNA damage (Fig. S6L). To test *in vivo* effects of aciclovir, mice bearing PC3 xenografts were treated with 25 mg valaciclovir per gram food, which gives aciclovir concentrations comparable to therapeutic concentrations in humans (Fig. S6M)^45,46^, in intervals of five days treatment and seven days intermission. Valaciclovir induced selective apoptosis in PC3 xenografts (Fig. S6N-O) and tumor growth was reduced by 16%, comparing mean volume of tumors at 6 weeks (Fig. 4D; p=0.018). Valaciclovir (n=16) increased survival (Fig. 4E; log-rank test; p=0.011) compared to control mice (n=15), with 31% (5/16) of animals alive 60 days after treatment initiation compared to 0% (0/15) of control (Fisher’s exact test p=0.043). In summary, the effect size of valaciclovir treatment on tumor size and survival of tumor bearing mice was modest but prompted us to examine its potential clinical use further.

We conducted a cohort study evaluating if men prescribed aciclovir had decreased risk of prostate cancer mortality using Danish population-based linked registries^47–50^. Since prostate cancer is common, slow growing and may be present years before diagnosis, we examined risk of prostate cancer mortality in all men aged 40 and older (Fig. 4F). Systemic aciclovir/valaciclovir users (from here defined as aciclovir users; n=156,116) were matched by age at first prescription, defined as start of follow-up, and calendar year to non-users (n=780,580; Fig. 4F, Table S1). All aciclovir-users, independent of the number of aciclovir prescriptions, were included. In cox proportional hazard models adjusted for age, calendar year and comorbidity index (Table S1), aciclovir users had no significant difference in risk of prostate cancer mortality, adjusted HR 0.95 (95% CI 0.90-1.01, p=0.09) (Fig. 4G). In conclusion, we do not find evidence to support general anti-cancer benefits of aciclovir in men at risk of prostate cancer mortality.

### Ellipticine and mithramycin A reduces CMV abundance and are effective in prostate cancer models

Ellipticine and mithramycin A reduces latent CMV genome replication^38^. CMV survival dependent cell lines (LNCaP, DU145) were more sensitive to ellipticine and mithramycin A compared to CMV survival independent cell lines (LAPC-4, PC3) (Fig. 5A). Both drugs reduced CMV DNA in DU145 cells (Fig. 5B). Ellipticine and mithramycin A acted independently of their known target genes TOPIIB and SP1, respectively, through which the drugs regulate latent CMV genome replication in hematopoietic cells^38^ (Fig. S7A-E). In light of these findings, we asked if mithramycin A and ellipticine reduced CMV genomes independent of cellular replication. LNCaP cells were continuously administered with EdU to label cells that undergo cell cycle S-phase. Ellipticine and mithramycin A reduced EdU incorporation (Fig. S7F) and induced apoptosis in both EdU^+^ and EdU^-^ cells (Fig. S7G). These results indicate that these drugs reduce, and possibly degrade, CMV independent of cellular replication. In contrast to ellipticine and mithramycin A, which can both act as DNA intercalators, two other chemotherapeutic drugs, cisplatin and etoposide, only readily reduced cell viability in DU145 cells (Fig. 5A) and etoposide did not reduce CMV DNA abundance (Fig. 5B).

**Figure 5:**
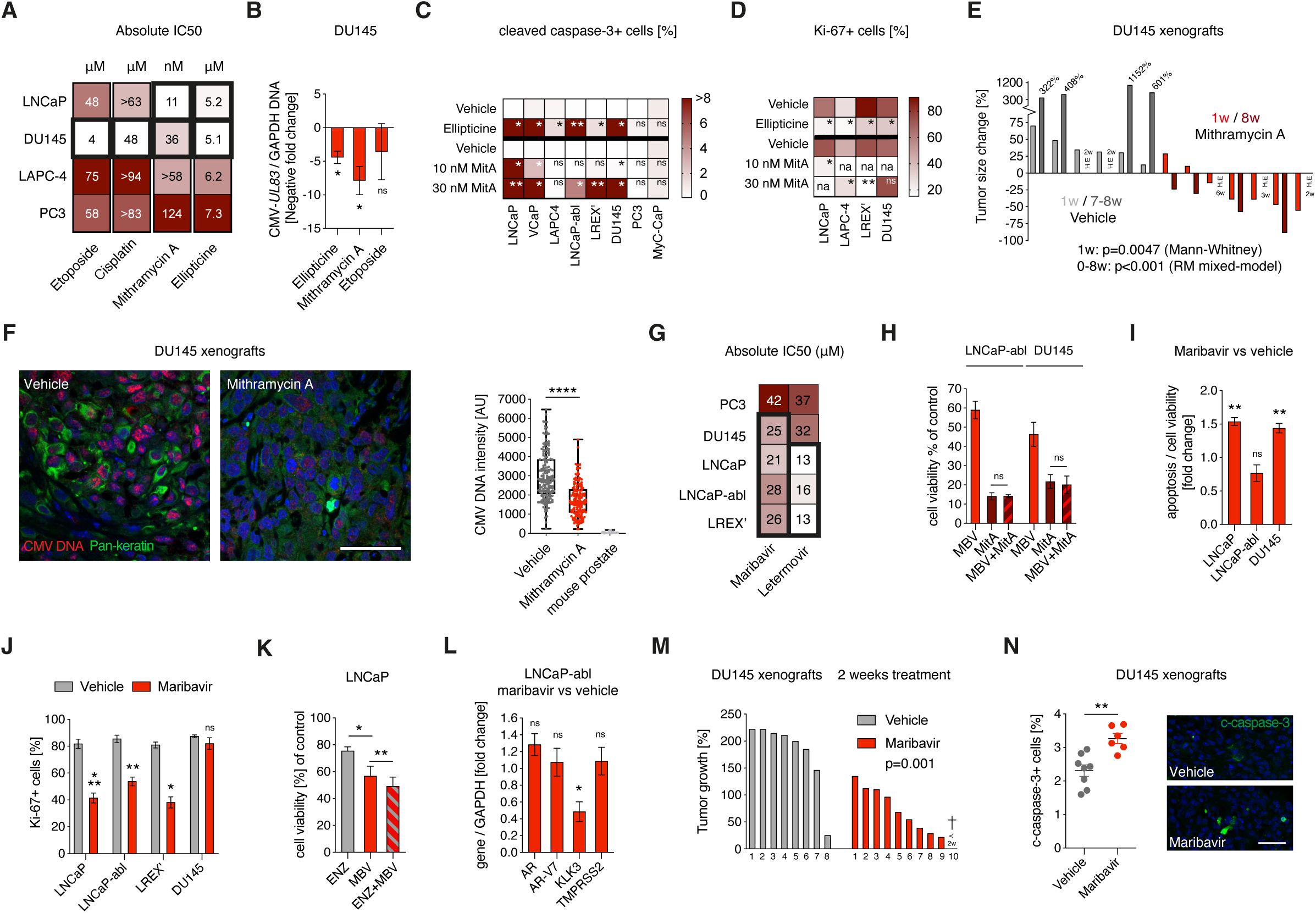
CMV can be therapeutically targeted in prostate cancer with chemotherapy and novel anti-viral drugs. **A)** Heat map with absolute IC50 in cell lines treated with cisplatin (n=2), etoposide (n=3), mithramycin A (n=4-5) or ellipticine (n=3-4) three days. Thick boxes show the most drug sensitive cell lines. **B)** CMV DNA abundance determined with CMV-*UL83* qPCR in DU145 four days after treatment with 3 μM ellipticine, 30 nM mithramycin A or 3 μM etoposide compared to vehicle (n=3). Data is displayed as negative fold change. **C)** Heat map showing mean of percentage cleaved caspase-3+ cells in cells treated with vehicle or 3 μM ellipticine (one day) and vehicle or 10-30 nM mithramycin A (three days) (n=3-5). **D)** Heat map showing mean of percentage Ki-67 in cells treated with vehicle or 3 μM ellipticine after one day and vehicle or 10-30 nM mithramycin A after three days (n=3-4). Na is not analyzed. **E)** Alteration of tumor size in percentage 1 week (w) and 7-8 weeks after vehicle (n=6) or mithramycin A treatment (n=7). 1w: Mann-Whitney t-test; 0-8w: Repeated Measures (RM) mixed-model. H.E = Humane endpoint (tumor ulcers: n=2 control, n=1 mithramycin A; found dead n=1 mithramycin A; unwell n=1 mithramycin A). **F)** CMV DNA examined by CMV FISH in tumors and mouse prostate after 7-8 weeks of treatment. Cell nuclei are labeled in blue with DAPI. Pan-keratin was used as a marker for epithelial cells. CMV DNA in pan-keratin+ cells, measured by nuclear intensity, was compared between vehicle (n=3) and mithramycin A (n=3) treated mice with un-paired t-test. Epithelial cells visualized with DAPI were quantified in mouse prostate. Scale bar: 50 μm. Box plot show median; box is 25-75 percentiles and error bars show max and min. **G)** Heat map showing absolute IC50 (μM) of letermovir and maribavir in cell lines, analyzed after six days of treatment (n=3-4). Thick boxes show the most drug sensitive cell lines. **H)** LNCaP-abl and DU145 were treated with 30 μM maribavir (MBV), 30 nM mithramycin A (MitA) or both. Cell viability was examined after three days of treatment (n=3). **I)** Cells were treated with vehicle or 30 μM maribavir and analyzed after three days for apoptosis, examined by the Caspase-Glo 3/7 assay, and cell viability (n=3-4). Apoptosis compared to total viable cells was represented as fold change. **J)** Cell lines treated with vehicle or 30 μM maribavir and analyzed after three days for percentage of Ki-67^+^ cells (n=3). **K)** Cell viability in LNCaP treated with 10 μM enzalutamide (ENZ), 30 μM maribavir (MBV) or both. (n=3). **L)** Gene expression assessed in LNCaP-abl after four days of treatment with maribavir compared to vehicle. Data is displayed as fold change. **M)** DU145 tumor growth in percentage after 2 weeks treatment with vehicle or 100 mg/kg maribavir two times per day. **N)** Percentage of cleaved caspase-3^+^ cells in vehicle (n=8) and maribavir (n=6) treated DU145 xenografts. Scale bar: 50 μm. Cell nuclei are labeled in blue with DAPI. Un-paired student’s t-test in (M)-(N). Paired t-tests were performed unless otherwise stated. Bar graphs are shown with mean value and error bars are SEM. P<0.05: *; p<0.01:**; p<0.001:***; p<0.0001:****, ns is non-significant.

Mithramycin A and ellipticine induced apoptosis in the same cell lines as IE1/2 RNAi (Fig. 5C) reduced proliferation (Fig. 5D) and AR target gene expression^51,52^. There were no additive effects of IE1/2 RNAi and ellipticine or mithramycin A on apoptosis or proliferation (Fig. S7H-I), suggesting that they all act through the same CMV dependent mechanism. In addition, mithramycin A reduced aciclovir-induced apoptosis in PC3 cells (Fig. S7J), further supporting a CMV driven response. Mithramycin A and ellipticine drastically reduce cell viability after three days (Fig. 5A), being both faster and more efficient than aciclovir, which did not alter cell viability in three days (Fig. S7K).

Mithramycin A is sometimes used to treat patients with testicular cancer^53^ and cancer-related hypercalcemia^54^, whereas ellipticine is not in clinical use. We asked whether Mithramycin A may affect prostate cancer cells in a xenograft model in mice. Mithramycin A induced shrinkage of DU145 cell xenografts after seven days of treatment that was durable over an 8-week period (Fig. 5E), displayed signs of active apoptosis (Fig. S7L)^55^ and reduced CMV genome abundance (Fig. 5F). Although mithramycin A can have severe side effects that limits its clinical utility, we conclude that CMV is an actionable target in CMV^+^ prostate cancer.

### Anti-cancer effects of novel anti-CMV compounds in models of prostate cancer

The CMV UL97 kinase inhibitor maribavir and the UL56 inhibitor letermovir are well-tolerated novel anti-viral drugs with *in vivo* anti-CMV efficiency in phase III clinical trials^56,57^. CMV dependent cell lines (CSPC, CRPC-Adeno and CRPC-SCL phenotypes) were sensitive to maribavir whereas letermovir sensitivity was limited to AR expressing cell lines (CSPC and CRPC-Adeno) (Fig. 5G, Fig. S8A), why we focused on maribavir. Co-treatment of maribavir with CMV reduction therapy had no additive effects on cell viability in LNCaP-abl and DU145 (Fig. 5H), indicating that maribavir acts through CMV. Maribavir promoted apoptosis in CSPC and CRPC-SCL cell lines (Fig. 5I) and reduced the percentage of proliferating Ki-67^+^ cells in CSPC and CRPC-Adeno cell lines (Fig. 5J). Similar to CMV reduction treatment such as mithramycin A, maribavir did not alter proliferation status in DU145 (Fig. 5J). Furthermore, PC3 was less sensitive to both maribavir and mithramycin A, compared to other cell lines which were more sensitive to both drugs (Fig. 5A, 5C-D, 5G). As for CMV depletion, maribavir had moderate additive effects to enzalutamide in LNCaP (Fig. 5K), suggesting that maribavir can inhibit cell viability through androgen signaling. Examined in LNCaP-abl, maribavir reduced expression of the AR target gene *KLK3* but did not alter expression of *AR*, *AR-V7* or *TMPRSS2* (Fig. 5L), showing that maribavir partly mimics CMV attenuation. Maribavir was nevertheless capable of inducing CMV dependent anti-cancer effects in CMV^+^ prostate cancer cell lines.

We examined *in vivo* effects of maribavir on prostate cancer growth. Mice with established DU145 xenografts were given 100 mg/kg maribavir (n=10) or vehicle (n=8) twice daily every weekday. Although animals lost weight the first week of maribavir treatment, they then stabilized and kept steady weight throughout the remaining treatment period (Fig. S8B), but continuous treatment for more than 3.5 weeks was not well tolerated. In line with the *in vitro* results, maribavir did not change the number of proliferating cells in DU145 xenografts (Fig. S8C). Notably, tumor growth was reduced in mice treated with maribavir examined after 2 weeks (Fig. 5M). Maribavir induced reduction of tumor growth was accompanied by increased apoptotic activity in the tumors (Fig. 5N). These results show that anti-CMV drugs have potential as a prostate cancer therapy.

## DISCUSSION

Here we report that latent CMV is commonly present in the healthy and malignant prostate epithelium. Endogenous latent CMV in prostate cancer cell lines was reduced in abundance by CMV-IE1/2 RNAi and by several pharmaceutical compounds. CMV promoted proliferation and cell survival in castration sensitive and resistant prostate cancer cell lines, partly via CMV-UL97 kinase and androgen signaling. Aciclovir showed modest effects on tumor growth in a pre-clinical model, but aciclovir usage in clinical practice was not associated with significantly reduced prostate cancer mortality at the dose and duration studied. Novel CMV drugs including the CMV-UL97 inhibitor maribavir had more marked effects on cell viability in CMV dependent prostate cancer cells. These results prompt investigation into the clinical use of anti-viral drugs such as maribavir for the treatment of prostate cancer.

The presence, characteristics and role of CMV in tissues and in diseases such as cancer is challenging to characterize^27^, not least due to inconsistencies in viral detection techniques. It is broadly accepted that cells of the hematopoietic lineage are infected by CMV, although CMV RNA is difficult to detect in chronic CMV infection^20,23,28,34^ and viral protein expression has rarely been assessed. Viral gene expression during latency may also be undetectable in HPV infected tumors even though classically viral-regulated genes can still be altered^58^. In hematopoietic cells, CMV may be easier to detect in CMV IgG^+^ individuals due to higher re-activation tendencies that triggers an anti-viral immune response. In 2003, Samanta et al., describe CMV infection in precancerous and cancerous epithelial cells in prostate biopsies from 22 patients, although no quantitative measures were reported^26^.

CMV persists in cells of the hematopoietic lineage as episomes^19^ via an interplay of viral and cellular proteins that allow replication of circular viral DNA^38^. The major IE locus (*UL122-UL123*) is not only an essential promotor of linear and episomal viral replication^37^, but our data suggest that latent CMV requires its expression to escape degradation, which was also induced by DNA intercalating chemotherapy. Perturbation of IE1/2 in endogenously CMV infected glioblastoma stem cells reduces cell viability^59^, perhaps pointing to shared functions of CMV in several tumor types. Crispr-Cas technology could potentially be used to remove latent viruses such as CMV^60^, but has so far only been applied on actively replicating CMV^61^. It is not known why gene expression is so difficult to detect during latent CMV infection, although still possible to functionally inhibit *in vitro*. CMV may have a complex RNA conformation which can act as an immune evasion mechanism. Multiple herpes viruses antagonize DNA sensing pathways, including STING-cGAS, to evade detection^62^ and latent hepatitis B virus can be degraded by induction of APOBEC cytidine deaminases^63^. The CMV proteins pp71 and pp65, both expressed in cells of prostate origin, can diminish an anti-viral response^64–66^. The tumor suppressor PTEN, that is deleted in around 20% of primary prostate cancers, promotes innate anti-viral sensing^67^. Interplays between cancer mutations and CMV infection may exist.

The majority of analyzed prostate cancer cell lines were dependent on endogenous CMV for cell survival, reminiscent of addiction to oncogenes such as MYC, a protein that, similar to latent viruses, promotes immune evasion^68^. CMV, with around 700 putative open reading frames^69^, can alter many cellular processes, including those that promote cell survival and proliferation, for viral replication and dissemination to proceed^70^. Latency may have other cellular consequences. It is therefore important to study viral effects on cancer in its proper context. In addition to cell survival, CMV promoted proliferation in part by influencing androgen signaling. Another potential mechanism could be through phosphorylation of Rb by UL97 (Fig. 2E)^36^. The percentage of cells in cell cycle was not altered by CMV inhibition, in Rb^low^ DU145, speaking for this hypothesis. Intertumoral heterogeneity in cellular consequences of CMV inhibition may be determined by different CMV strains. Importantly, several CMV inhibition methods reduced cell viability in CRPC cells, a disease state with limited treatment options. A multitude of resistance mechanisms have been described, including AR mutations, *AR* or *AR-V7* upregulation and lineage switching from adenocarcinoma to androgen indifferent basal/neuroendocrine cell states^40^. CMV may potentially cause or exacerbate anti-androgen resistance.

Although aciclovir induced CMV dependent DNA damage and apoptosis *in vitro*, the *in vivo* effect was modest, and we found no association between repeated aciclovir usage and prostate cancer mortality. Although these results speak against clinical activity of aciclovir, identifying possible super responders, exploring timing and duration of treatment, dosing schemes and combination therapies will be important to further evaluate possible clinical benefits. The novel anti-viral drugs letermovir and maribavir had more substantial effects on prostate cancer cell viability than aciclovir at therapeutic doses, with maribavir having the broadest effects. Mithramycin A was most efficient in mimicking experimental CMV loss, but systemic toxicities hamper general clinical use. Milder mithramycin A analogues and novel drug delivery systems could potentially be used to overcome this obstacle. CMV abundance, functions of CMV and the application of anti-viral drugs in other tumor types are interesting avenues to explore.

## ACKNOWLEDGEMENTS

We thank members of the Frisén lab for discussions, especially Camilla Engblom and Jeff Mold. We thank Sarantis Giatrellis for assistance with FACS. We thank Dr. Olof Breuer for consulting on aciclovir pharmacology applied to mouse studies. We thank Cecilia Söderberg-Nauclér and members of her laboratory for fruitful discussions and support in the initiation of this study. We thank Maria Genander and Anna Mourskaia for valuable input on animal study design and execution.

This study was supported by grants from the Swedish Research Council (D0761801), the Swedish Cancer Society (19 0452 Pj 01 H), the Swedish Foundation for Strategic Research (SB16-0014) and Knut och Alice Wallenbergs Stiftelse (2018.0063). The PCBN biobank is supported by the Department of Defense Prostate Cancer Research Program Award No W81XWH-14-2-0182, W81XWH-14-2-0183, W81XWH-14-2-0185, W81XWH-14-2-0186, and W81XWH-15-2-0062 Prostate Cancer Biorepository Network. For CPC-GENE data, the computations and data handling were enabled by resources provided by the Swedish National Infrastructure for Computing (SNIC) at UPPMAX partially funded by the Swedish Research Council through grand agreement no. 2018-05973. We thank the scientific teams of CPC-GENE for allowing us to analyze their cohort through the International Cancer Genome Consortium. J.C and M.S were supported by the Clinical Scientist Training Program at Karolinska Institutet.

## AUTHOR CONTRIBUTIONS

J.F. and J.C. conceived the study. J.F. supervised the study. J.C. performed cell experiments, analysis of human tissue and blood. J.C., M.S., M.Z., C-J.E. and A.S. performed and analyzed animal experiments. K.A. and H.D. procured tissue and blood from men post-mortem. K.A. collected post-mortem tissues and blood and stained FFPE sections with H&E. H.D. performed histological assessments of prostate tissue and reviewed medical history. A.T. and A.S. collected data on human samples. P.W. designed collection of bone metastases and provided tissues for analysis. L.P., H-O.A, H.T.S. designed epidemiological studies. L.P. performed statistical analysis of epidemiological studies. J.F. and J.C. wrote the manuscript with intellectual input from all authors.

## SUPPLEMENTARY INFORMATION

**Supplemental Figure 1-8**

**Supplemental Table 1**

**Online Methods**

**Supplemental References**

## Notes

### Competing Interest Statement

The authors have declared no competing interest.

## REFERENCES

1. Sandhu, S., et al. Prostate cancer. Lancet 398, 1075–1090 (2021).

2. Virgin, H.W., Wherry, E.J. & Ahmed, R. Redefining chronic viral infection. Cell 138, 30–50 (2009).

3. Virgin, H.W. The virome in mammalian physiology and disease. Cell 157, 142–150 (2014).

4. Ang, K.K., et al. Human papillomavirus and survival of patients with oropharyngeal cancer. N Engl J Med 363, 24–35 (2010).

5. McBride, A.A. Human papillomaviruses: diversity, infection and host interactions. Nat Rev Microbiol 20, 95–108 (2022).

6. Zuhair, M., et al. Estimation of the worldwide seroprevalence of cytomegalovirus: A systematic review and meta-analysis. Rev Med Virol 29, e2034 (2019).

7. Stanier, P., et al. Persistence of cytomegalovirus in mononuclear cells in peripheral blood from blood donors. BMJ 299, 897–898 (1989).

8. Taylor-Wiedeman, J., Sissons, J.G., Borysiewicz, L.K. & Sinclair, J.H. Monocytes are a major site of persistence of human cytomegalovirus in peripheral blood mononuclear cells. J Gen Virol 72 (Pt 9), 2059–2064 (1991).

9. Kraat, Y.J., et al. Detection of latent human cytomegalovirus in organ tissue and the correlation with serological status. Transpl Int 5 **Suppl 1**, S613–616 (1992).

10. Zhang, L.J., Hanff, P., Rutherford, C., Churchill, W.H. & Crumpacker, C.S. Detection of human cytomegalovirus DNA, RNA, and antibody in normal donor blood. J Infect Dis 171, 1002–1006 (1995).

11. Mendelson, M., Monard, S., Sissons, P. & Sinclair, J. Detection of endogenous human cytomegalovirus in CD34+ bone marrow progenitors. J Gen Virol 77 **(Pt** **12**), 3099–3102 (1996).

12. Hendrix, R.M., Wagenaar, M., Slobbe, R.L. & Bruggeman, C.A. Widespread presence of cytomegalovirus DNA in tissues of healthy trauma victims. J Clin Pathol 50, 59–63 (1997).

13. Larsson, S., Soderberg-Naucler, C., Wang, F.Z. & Moller, E. Cytomegalovirus DNA can be detected in peripheral blood mononuclear cells from all seropositive and most seronegative healthy blood donors over time. Transfusion 38, 271–278 (1998).

14. Roback, J.D., et al. Multicenter evaluation of PCR methods for detecting CMV DNA in blood donors. Transfusion 41, 1249–1257 (2001).

15. Okedele, O.O., Nelson, H.H., Oyenuga, M.L., Thyagarajan, B. & Prizment, A. Cytomegalovirus and cancer-related mortality in the national health and nutritional examination survey. Cancer Causes Control 31, 541–547 (2020).

16. Sutcliffe, S., et al. Prospective study of cytomegalovirus serostatus and prostate cancer risk in the Prostate Cancer Prevention Trial. Cancer Causes Control 23, 1511–1518 (2012).

17. Classon J, Britten A, Alkass K, Druid H, Brenner N, Waterboer T, Wareham N J, Gkrania-Klotsas E, Frisén J The role of cytomegalovirus in prostate cancer incidence and mortality. BioRxiv (2023).

18. Classon, J., et al. Prostate cancer disease recurrence after radical prostatectomy is associated with HLA type and local cytomegalovirus immunity. Mol Oncol (2022).

19. Bolovan-Fritts, C.A., Mocarski, E.S. & Wiedeman, J.A. Peripheral blood CD14(+) cells from healthy subjects carry a circular conformation of latent cytomegalovirus genome. Blood 93, 394–398 (1999).

20. Sinclair, J. & Sissons, P. Latency and reactivation of human cytomegalovirus. J Gen Virol 87, 1763–1779 (2006).

21. Nikolich-Zugich, J., Goodrum, F., Knox, K. & Smithey, M.J. Known unknowns: how might the persistent herpesvirome shape immunity and aging? Curr Opin Immunol 48, 23–30 (2017).

22. Gordon, C.L., et al. Tissue reservoirs of antiviral T cell immunity in persistent human CMV infection. J Exp Med 214, 651–667 (2017).

23. Shnayder, M., et al. Defining the Transcriptional Landscape during Cytomegalovirus Latency with Single-Cell RNA Sequencing. mBio 9 (2018).

24. Boldogh, I., Baskar, J.F., Mar, E.C. & Huang, E.S. Human cytomegalovirus and herpes simplex type 2 virus in normal and adenocarcinomatous prostate glands. J Natl Cancer Inst 70, 819–826 (1983).

25. Kuhn, J.E., et al. Quantitation of human cytomegalovirus genomes in the brain of AIDS patients. J Med Virol 47, 70–82 (1995).

26. Samanta, M., Harkins, L., Klemm, K., Britt, W.J. & Cobbs, C.S. High prevalence of human cytomegalovirus in prostatic intraepithelial neoplasia and prostatic carcinoma. J Urol 170, 998–1002 (2003).

27. Wick, W. & Platten, M. CMV infection and glioma, a highly controversial concept struggling in the clinical arena. Neuro Oncol 16, 332–333 (2014).

28. Cheng, S., et al. Transcriptome-wide characterization of human cytomegalovirus in natural infection and experimental latency. Proc Natl Acad Sci U S A 114, E10586–E10595 (2017).

29. Tang, K.W., Alaei-Mahabadi, B., Samuelsson, T., Lindh, M. & Larsson, E. The landscape of viral expression and host gene fusion and adaptation in human cancer. Nat Commun 4, 2513 (2013).

30. Zapatka, M., et al. The landscape of viral associations in human cancers. Nat Genet 52, 320–330 (2020).

31. Jackson, S.E., et al. Latent Cytomegalovirus (CMV) Infection Does Not Detrimentally Alter T Cell Responses in the Healthy Old, But Increased Latent CMV Carriage Is Related to Expanded CMV-Specific T Cells. Front Immunol 8, 733 (2017).

32. Parry, H.M., et al. Cytomegalovirus viral load within blood increases markedly in healthy people over the age of 70 years. Immun Ageing 13, 1 (2016).

33. Slobedman, B. & Mocarski, E.S. Quantitative analysis of latent human cytomegalovirus. J Virol 73, 4806–4812 (1999).

34. Roback, J.D., et al. CMV DNA is rarely detected in healthy blood donors using validated PCR assays. Transfusion 43, 314–321 (2003).

35. Thysell, E., et al. Gene expression profiles define molecular subtypes of prostate cancer bone metastases with different outcomes and morphology traceable back to the primary tumor. Mol Oncol 13, 1763–1777 (2019).

36. Hume, A.J., et al. Phosphorylation of retinoblastoma protein by viral protein with cyclin-dependent kinase function. Science 320, 797–799 (2008).

37. Collins-McMillen, D., Buehler, J., Peppenelli, M. & Goodrum, F. Molecular Determinants and the Regulation of Human Cytomegalovirus Latency and Reactivation. Viruses 10 (2018).

38. Tarrant-Elorza, M., Rossetto, C.C. & Pari, G.S. Maintenance and replication of the human cytomegalovirus genome during latency. Cell Host Microbe 16, 43–54 (2014).

39. Tang, F., et al. Chromatin profiles classify castration-resistant prostate cancers suggesting therapeutic targets. Science 376, eabe1505 (2022).

40. Watson, P.A., Arora, V.K. & Sawyers, C.L. Emerging mechanisms of resistance to androgen receptor inhibitors in prostate cancer. Nat Rev Cancer 15, 701–711 (2015).

41. Jurhill, R.R., van der Veen, H., van Leenders, G.J. & Verhagen, P.C. Reduction of serum prostate-specific antigen levels following varicella-zoster infection and valaciclovir treatment in prostate cancer. Eur Urol 56, 392–394 (2009).

42. Talarico, C.L., et al. Acyclovir is phosphorylated by the human cytomegalovirus UL97 protein. Antimicrob Agents Chemother 43, 1941–1946 (1999).

43. Littler, E., Stuart, A.D. & Chee, M.S. Human cytomegalovirus UL97 open reading frame encodes a protein that phosphorylates the antiviral nucleoside analogue ganciclovir. Nature 358, 160–162 (1992).

44. Furman, P.A., St Clair, M.H. & Spector, T. Acyclovir triphosphate is a suicide inactivator of the herpes simplex virus DNA polymerase. J Biol Chem 259, 9575–9579 (1984).

45. Weller, S., et al. Pharmacokinetics of the acyclovir pro-drug valaciclovir after escalating single-and multiple-dose administration to normal volunteers. Clin Pharmacol Ther 54, 595–605 (1993).

46. Blum, M.R., Liao, S.H. & de Miranda, P. Overview of acyclovir pharmacokinetic disposition in adults and children. Am J Med 73, 186–192 (1982).

47. Gjerstorff, M.L. The Danish Cancer Registry. Scand J Public Health 39, 42–45 (2011).

48. Pottegard, A., et al. Data Resource Profile: The Danish National Prescription Registry. Int J Epidemiol 46, 798–798f (2017).

49. Schmidt, M., Pedersen, L. & Sorensen, H.T. The Danish Civil Registration System as a tool in epidemiology. Eur J Epidemiol 29, 541–549 (2014).

50. Schmidt, M., et al. The Danish National Patient Registry: a review of content, data quality, and research potential. Clin Epidemiol 7, 449–490 (2015).

51. Haffner, M.C., et al. Androgen-induced TOP2B-mediated double-strand breaks and prostate cancer gene rearrangements. Nat Genet 42, 668–675 (2010).

52. Wang, L.G. & Ferrari, A.C. Mithramycin targets sp1 and the androgen receptor transcription level-potential therapeutic role in advanced prostate cancer. Transl Oncogenomics 1, 19–31 (2006).

53. Brown, J.H. & Kennedy, B.J. Mithramycin in the Treatment of Disseminated Testicular Neoplasms. N Engl J Med 272, 111–118 (1965).

54. Singer, F.R., et al. Mithramycin treatment of intractable hypercalcemia due to parathyroid carcinoma. N Engl J Med 283, 634–636 (1970).

55. Choi, E.S., et al. Myeloid cell leukemia-1 is a key molecular target for mithramycin A-induced apoptosis in androgen-independent prostate cancer cells and a tumor xenograft animal model. Cancer Lett 328, 65–72 (2013).

56. Maertens, J., et al. Maribavir for Preemptive Treatment of Cytomegalovirus Reactivation. N Engl J Med 381, 1136–1147 (2019).

57. Marty, F.M., et al. Letermovir Prophylaxis for Cytomegalovirus in Hematopoietic-Cell Transplantation. N Engl J Med 377, 2433–2444 (2017).

58. Puram, S.V., et al. Cellular states are coupled to genomic and viral heterogeneity in HPV-related oropharyngeal carcinoma. Nat Genet 55, 640–650 (2023).

59. Soroceanu, L., et al. Cytomegalovirus Immediate-Early Proteins Promote Stemness Properties in Glioblastoma. Cancer Res 75, 3065–3076 (2015).

60. White, M.K., Hu, W. & Khalili, K. The CRISPR/Cas9 genome editing methodology as a weapon against human viruses. Discov Med 19, 255–262 (2015).

61. Hein, M.Y. & Weissman, J.S. Functional single-cell genomics of human cytomegalovirus infection. Nat Biotechnol (2021).

62. Paludan, S.R. & Bowie, A.G. Immune sensing of DNA. Immunity 38, 870–880 (2013).

63. Lucifora, J., et al. Specific and nonhepatotoxic degradation of nuclear hepatitis B virus cccDNA. Science 343, 1221–1228 (2014).

64. Trgovcich, J., Cebulla, C., Zimmerman, P. & Sedmak, D.D. Human cytomegalovirus protein pp71 disrupts major histocompatibility complex class I cell surface expression. J Virol 80, 951–963 (2006).

65. Fu, Y.Z., et al. Human Cytomegalovirus Tegument Protein UL82 Inhibits STING-Mediated Signaling to Evade Antiviral Immunity. Cell Host Microbe 21, 231–243 (2017).

66. Browne, E.P. & Shenk, T. Human cytomegalovirus UL83-coded pp65 virion protein inhibits antiviral gene expression in infected cells. Proc Natl Acad Sci U S A 100, 11439–11444 (2003).

67. Li, S., et al. The tumor suppressor PTEN has a critical role in antiviral innate immunity. Nat Immunol 17, 241–249 (2016).

68. Dhanasekaran, R., et al. The MYC oncogene -the grand orchestrator of cancer growth and immune evasion. Nat Rev Clin Oncol (2021).

69. Stern-Ginossar, N., et al. Decoding human cytomegalovirus. Science 338, 1088–1093 (2012).

70. Cobbs, C. Cytomegalovirus is a tumor-associated virus: armed and dangerous. Curr Opin Virol 39, 49–59 (2019).

71. Webel, R., et al. Differential properties of cytomegalovirus pUL97 kinase isoforms affect viral replication and maribavir susceptibility. J Virol 88, 4776–4785 (2014).

