## Supplemental Information for "Cytomegalovirus promotes proliferation and survival of prostate cancer cells and constitutes a therapeutic target"

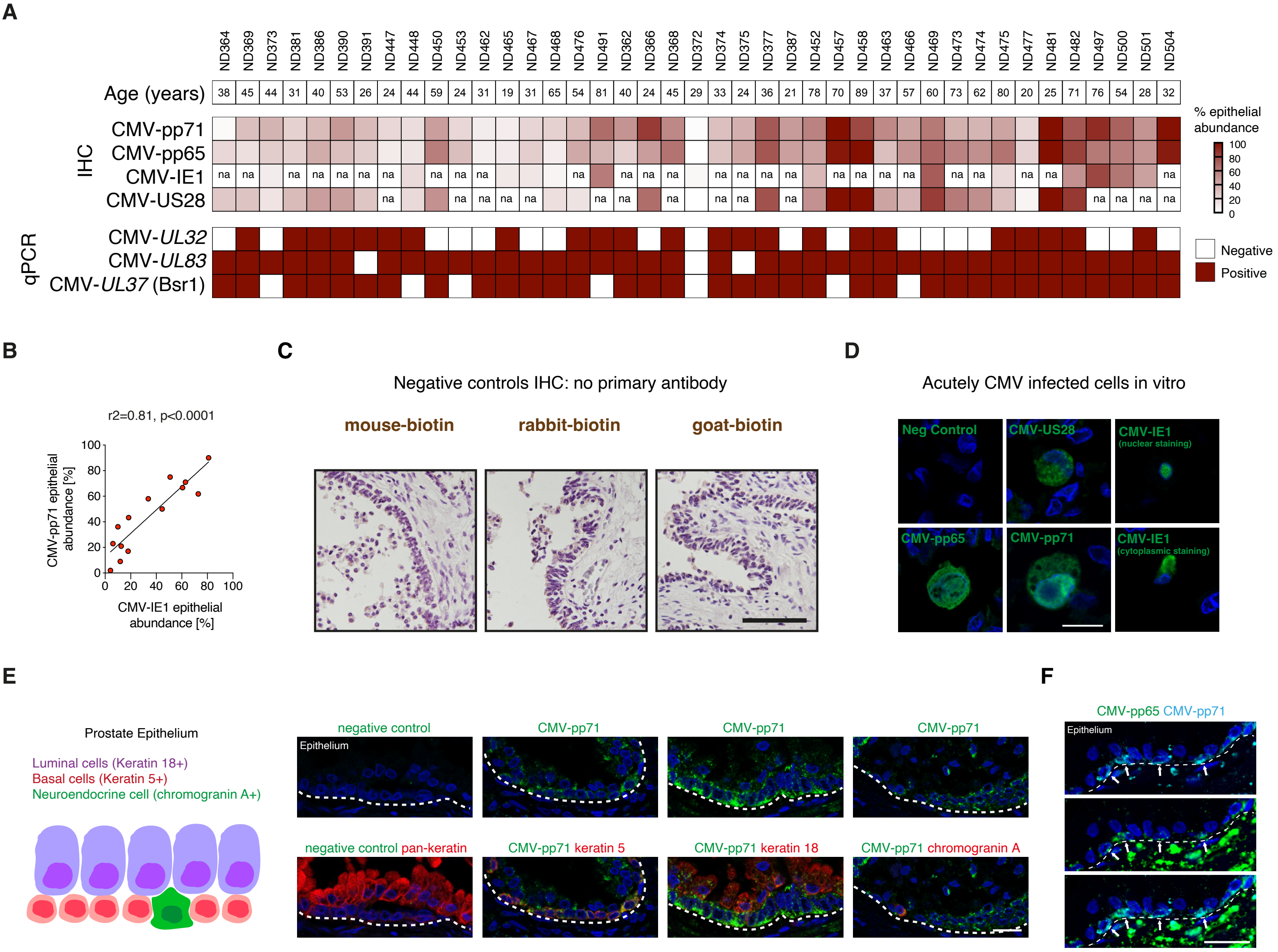
A
ND364 ND369 ND373 ND381 ND386 ND390 ND391 ND447 ND448 ND450 ND453 ND462 ND465 ND467 ND468 ND476 ND491 ND362 ND366 ND368 ND372 ND374 ND375 ND377 ND387 ND452 ND457 ND458 ND463 ND466 ND469 ND473 ND474 ND475 ND477 ND481 ND482 ND497 ND500 ND501 ND504
Age (years)
38 45 44 31 40 53 26 24 44 59 24 31 19 31 65 54 81 40 24 45 29 33 24 36 21 78 70 89 37 57 60 73 62 80 20 25 71 76 54 28 32
IHC
CMV-pp71
CMV-pp65
CMV-IE1
CMV-US28
na
% epithelial abundance
100
80
60
40
20
0
qPCR
CMV-*UL32*
CMV-*UL83*
CMV-*UL37* (Bsr1)
Negative
Positive
B
r2=0.81, p<0.0001
CMV-pp71 epithelial abundance [%]
CMV-IE1 epithelial abundance [%]
0 20 40 60 80 100
C
Negative controls IHC: no primary antibody
mouse-biotin
rabbit-biotin
goat-biotin
D
Acutely CMV infected cells in vitro
Neg Control
CMV-US28
CMV-IE1 (nuclear staining)
CMV-pp65
CMV-pp71
CMV-IE1 (cytoplasmic staining)
E
Prostate Epithelium
Luminal cells (Keratin 18+)
Basal cells (Keratin 5+)
Neuroendocrine cell (chromogranin A+)
negative control
Epithelium
CMV-pp71
CMV-pp71
CMV-pp71
negative control pan-keratin
CMV-pp71 keratin 5
CMV-pp71 keratin 18
CMV-pp71 chromogranin A
F
CMV-pp65 CMV-pp71
Epithelium

**Supplemental Figure 1: CMV DNA and protein in the prostate**

**A)** Heat map to summarize CMV detection in postmortem donors. CMV epithelial abundance was analyzed with four CMV IHC assays detecting CMV-pp71, CMV-pp65, CMV-IE1 and CMV-US28 respectively. Heat map shows epithelial abundance of CMV in percentage. CMV DNA was examined with three CMV qPCR assays; CMV-UL32, CMV-UL83 and CMV-UL37 (DNA pre-treated with *bsr1*). Na is not analyzed. **B)** Linear regression comparing CMV-pp71 epithelial abundance with CMV-IE1 epithelial abundance in prostate in percentage. **C)** CMV IHC assays with no primary antibody resulted in no DAB detection in prostate tissue. Three different secondary antibodies were used: mouse-biotin, rabbit-biotin or goat-biotin. Cell nuclei are labeled with hematoxylin in purple. Scale bar: 50 μm. **D)** IHC of acutely CMV infected cells *in vitro.* No staining was observed when primary antibody was omitted in negative (neg) control. CMV proteins were detected in cytoplasm and cell nuclei. Scale bar: 20 μm. Cell nuclei are labeled in blue with DAPI. **E)** Illustration and staining of CMV-pp71 in three main cell types of the prostate epithelium; basal cells (Keratin $5^{+}$), luminal cells (keratin $18^{+}$), neuroendocrine cells (chromogranin $A^{+}$). Pan-keratin was used as a marker for all epithelial cells. Scale bar: 20 μm. **F)** Co-labeling of CMV-pp65 (green) and CMV-pp71 (magenta). Arrows point to examples of epithelial cells positive for both proteins. Scale bar: 25 μm. Cell nuclei are labeled in blue with DAPI. Dotted line in (E) and (F) depicts the basal lamina of the epithelium.

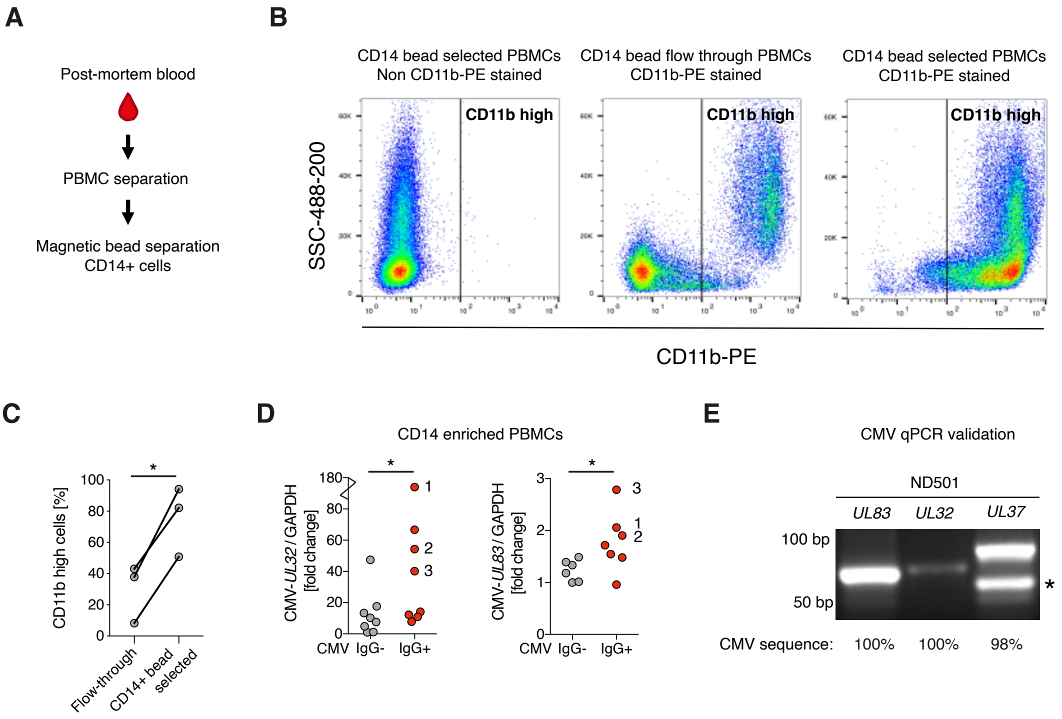
A
Post-mortem blood
PBMC separation
Magnetic bead separation
CD14+ cells
B
CD14 bead selected PBMCs
Non CD11b-PE stained
CD14 bead flow through PBMCs
CD11b-PE stained
CD14 bead selected PBMCs
CD11b-PE stained
CD11b high
SSC-488-200
CD11b-PE
C
CD11b high cells [%]
Flow-through
CD14+ bead selected
D
CD14 enriched PBMCs
CMV-UL32 / GAPDH [fold change]
CMV-UL83 / GAPDH [fold change]
CMV IgG- IgG+
E
CMV qPCR validation
ND501
UL83
UL32
UL37
100 bp
50 bp
CMV sequence: 100% 100% 98%

**Supplemental Figure 2: Validation of CMV qPCR**

**A)** Illustration of $CD14^+$ cell enrichment from blood in post-mortem donors. **B)** Bead enrichment of $CD14^+$ cells (which include monocytes) from peripheral blood mononuclear cells (PBMC) was validated with FACS. Cells were stained with a PE conjugated antibody against CD11b, which labels myeloid lineage cells. Unstained CD14 enriched cells (panel 1), CD11b stained CD14 bead flow through (panel 2) and CD11b stained CD14 enriched cells (panel 3) were examined. FACS plots show side scatter (SSC) on y-axes and CD11b-PE intensity on x-axes. The highest density of cells is shown in red, with decreasing density in yellow, green and blue. **C)** Percentage of CD11b high cells in CD11b stained CD14 bead flow through and CD11b stained CD14 enriched cells (n=3) were compared with paired t-test. **D)** qPCR of CMV-*UL32* and CMV-*UL83* in CD14 enriched PBMC DNA comparing CMV $IgG^+$ (n=8) and CMV $IgG^-$ donors (n=8) with un-paired two-sided t-test. Data points 1, 2 and 3 are labeled in the two graphs for comparison. **E)** Representative gel of prostate qPCR products with taqman primer/probes UL83, UL32 and UL37. For UL37 qPCR DNA was pre-treated with restriction enzyme bsr1. Sanger sequencing of cloned PCR products showed 100%, 100% and 98% match respectively to the merlin CMV genome. Asterisk point to CMV *UL37* PCR product. Bp = base pairs. $P<0.05$: *

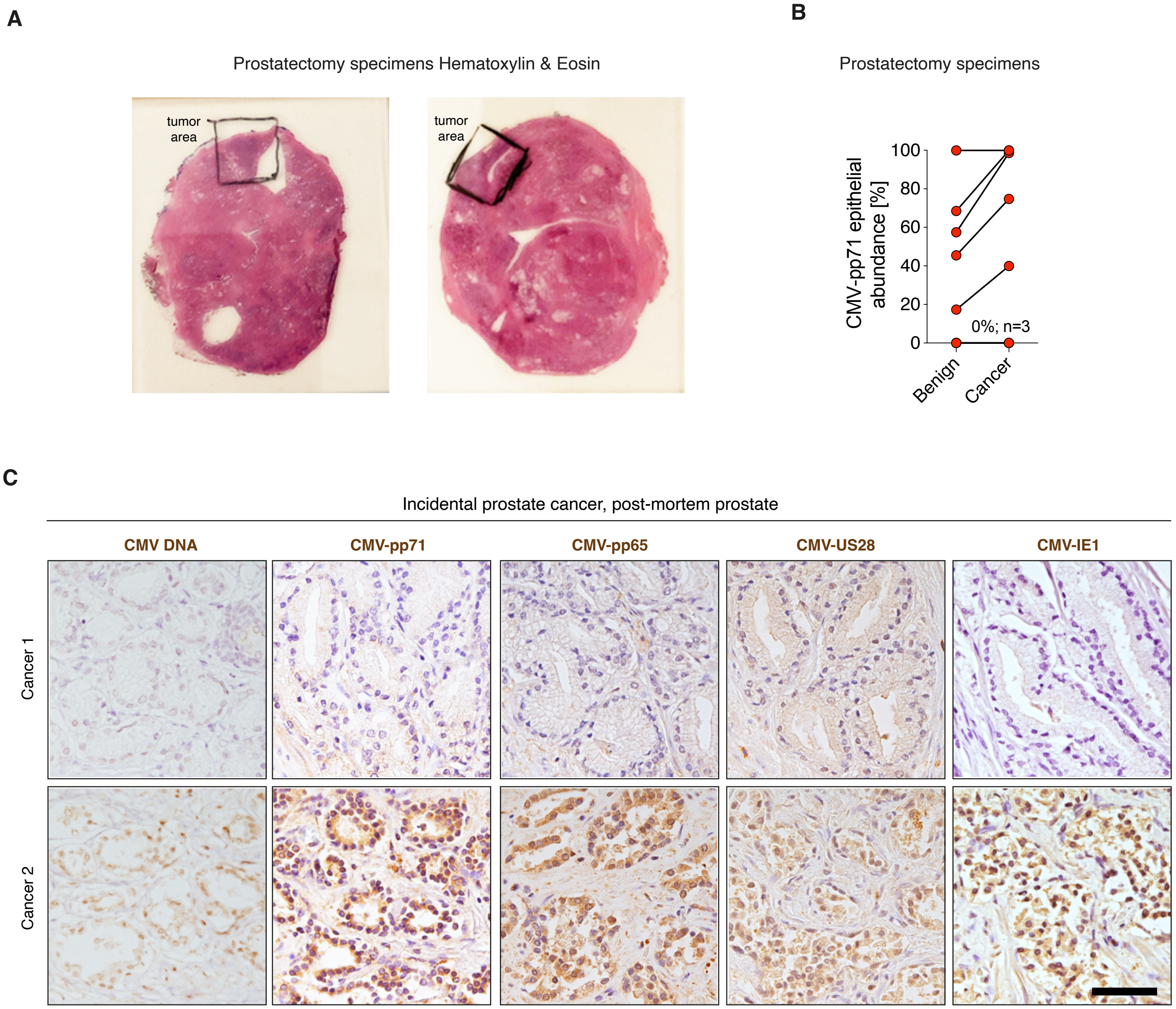
A
Prostatectomy specimens Hematoxylin & Eosin
tumor area
tumor area
B
Prostatectomy specimens
CMV-pp71 epithelial abundance [%]
100
80
60
40
20
0
0%; n=3
Benign
Cancer
C
Incidental prostate cancer, post-mortem prostate
CMV DNA
CMV-pp71
CMV-pp65
CMV-US28
CMV-IE1
Cancer 1
Cancer 2

**Supplemental Figure 3: CMV in prostate cancer**

**A)** H&E of prostatectomy specimens. Scale bar: 1 cm. Tumor areas were annotated by a pathologist associated with PCBN. **B)** CMV-pp71 abundance (%) in matched benign and cancer epithelium in prostatectomy specimens (n=8). **C)** Prostate cancer in one post-mortem donor (Cancer 1) was negative and suspected prostate cancer in another post-mortem donor was positive (Cancer 2) for CMV by in situ hybridization (CMV DNA) and CMV IHC. Cell nuclei are labeled with hematoxylin. Scale bar: 100 μm.

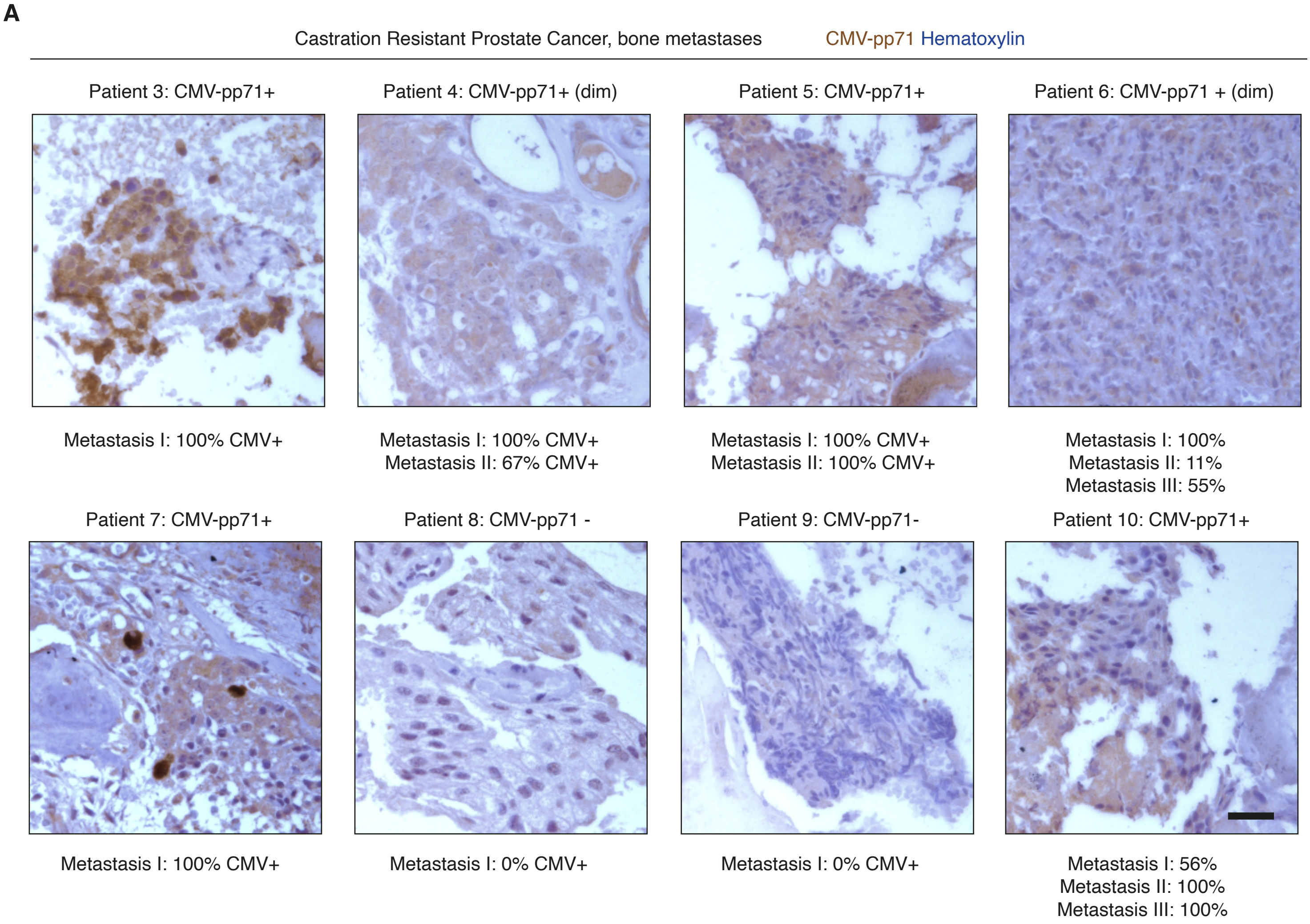
A
Castration Resistant Prostate Cancer, bone metastases
CMV-pp71 Hematoxylin
Patient 3: CMV-pp71+
Patient 4: CMV-pp71+ (dim)
Patient 5: CMV-pp71+
Patient 6: CMV-pp71 + (dim)
Metastasis I: 100% CMV+
Metastasis I: 100% CMV+
Metastasis II: 67% CMV+
Metastasis I: 100% CMV+
Metastasis II: 100% CMV+
Metastasis I: 100%
Metastasis II: 11%
Metastasis III: 55%
Patient 7: CMV-pp71+
Patient 8: CMV-pp71 -
Patient 9: CMV-pp71-
Patient 10: CMV-pp71+
Metastasis I: 100% CMV+
Metastasis I: 0% CMV+
Metastasis I: 0% CMV+
Metastasis I: 56%
Metastasis II: 100%
Metastasis III: 100%

**Supplemental Figure 4: CMV in prostate cancer metastases**

Representative images of CMV-pp71 IHC in CRPC bone metastases. In five patients, more than one metastasis was examined, labelled metastases I, II and III and percentage of CMV-pp71+ areas are shown in figure. Brown is CMV-pp71 and violet is hematoxylin staining that label cell nuclei. Scale bar: 50 μm.

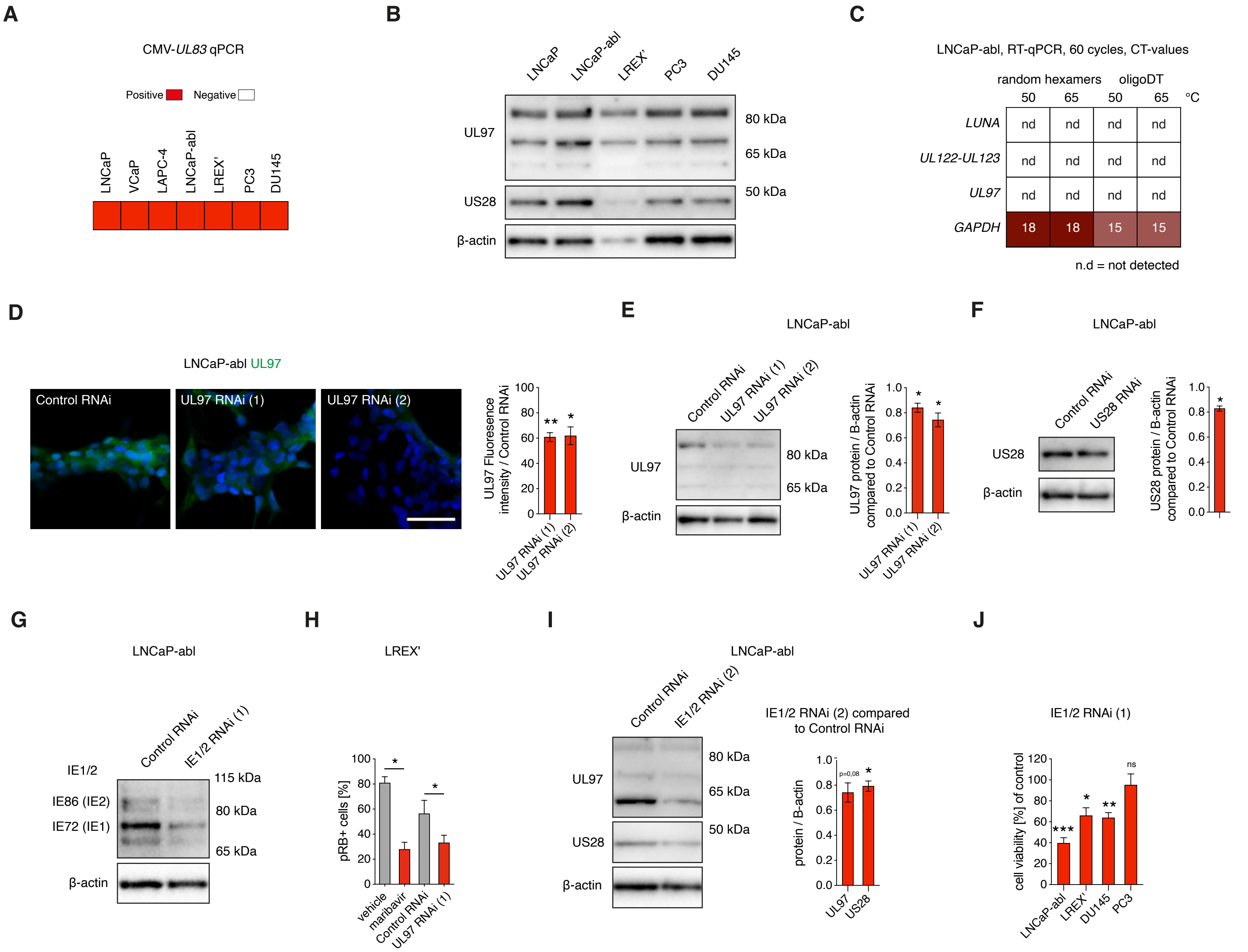
A
CMV-UL83 qPCR
Positive
Negative
LNCaP
VCaP
LAPC-4
LNCaP-abl
LREX'
PC3
DU145
B
LNCaP
LNCaP-abl
LREX'
PC3
DU145
UL97
US28
β-actin
80 kDa
65 kDa
50 kDa
C
LNCaP-abl, RT-qPCR, 60 cycles, CT-values
random hexamers
oligoDT
50
65
50
65
°C
LUNA
UL122-UL123
UL97
GAPDH
nd
18
18
15
15
n.d = not detected
D
LNCaP-abl UL97
Control RNAi
UL97 RNAi (1)
UL97 RNAi (2)
UL97 Fluoresence intensity / Control RNAi
E
LNCaP-abl
UL97 protein / B-actin compared to Control RNAi
F
LNCaP-abl
US28 RNAi
US28 protein / B-actin compared to Control RNAi
G
LNCaP-abl
IE1/2
IE86 (IE2)
IE72 (IE1)
IE1/2 RNAi (1)
115 kDa
H
LREX'
pRB+ cells [%]
vehicle
maribavir
I
IE1/2 RNAi (2)
IE1/2 RNAi (2) compared to Control RNAi
protein / B-actin
p=0,08
J
IE1/2 RNAi (1)
cell viability [%] of control
ns

**Supplemental Figure 5: CMV proteins are detected in prostate cancer cells and can be reduced with RNAi**

**A)** qPCR of CMV-*UL83* in cell lines. **B)** Immunoblot of cell lines. CMV-US28, CMV-UL97 and β-actin expression were examined. **C)** Expression of the CMV genes *LUNA, UL122-UL123* and *UL97* was not detected in LNCaP-abl, independent of cDNA synthesis protocol (random hexamers or oligoDT) or cDNA synthesis temperature (50 or 65 $^{o}$C). nd is not detected. **D)** Mean UL97 fluorescence intensity in LNCaP-abl after three days transfection with control RNAi or UL97 RNAi (1). Scale bar 50 μm. Cell nuclei are labeled in blue with DAPI. **E)** Immunoblot of LNCaP-abl four days after transfection with control RNAi, UL97 RNAi (1) or UL97 RNAi (2). CMV-UL97 and β-actin expression was examined and quantified (n=3). **F)** Immunoblot of LNCaP-abl four days after transfection with control RNAi or US28 RNAi. CMV-US28 and β-actin expression was examined and quantified (n=3). **G)** Immunoblot of LNCaP-abl treated four days with control RNAi or IE1/2 RNAi (1). Note that the classical IE isoforms IE72 (IE1) and IE86 (IE2) were present and downregulated with IE1/2 RNAi (1) treatment. Experiment was performed in replicate. **H)** Percentage of $pRB^+$ cells in LREX' three days after treatment with control RNAi or UL97 RNAi (1) and vehicle or 10 μM maribavir (n=3). **I)** Immunoblot of LNCaP-abl four days after transfection with control RNAi or IE1/2 RNAi (2). CMV-US28, CMV-UL97 and β-actin expression was examined and quantified (n=3). **J)** Cell viability in % compared to control, four days after treatment with control RNAi and IE1/2 RNAi (1) (n=3-5). Data in bar graphs are shown as mean. Error bars in bar graphs are SEM. Treatments were compared with paired t-tests in (H) and with one-sample t-test in (D), (E), (F), (I) and (J). Cell nuclei are labeled in blue with DAPI. $p<0.05$ is*, $p<0.01$ is **, $p<0.001$ is ***. Ns is non-significant. kDa is kilodalton.

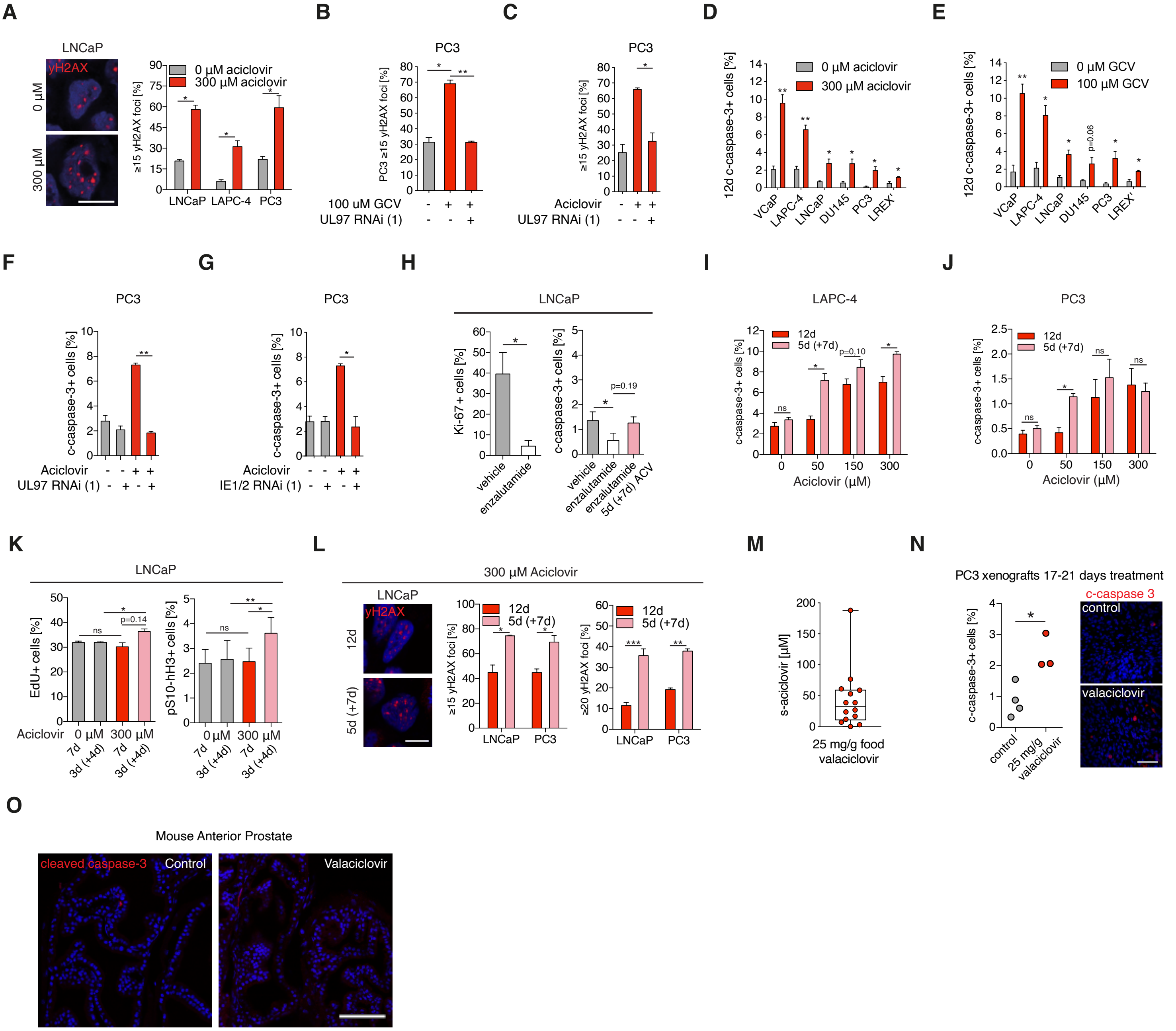
A
LNCaP
yH2AX
0 µM
300 µM
0 µM aciclovir
300 µM aciclovir
≥15 yH2AX foci [%]
LNCaP LAPC-4 PC3
B
PC3
PC3 ≥15 yH2AX foci [%]
100 uM GCV
UL97 RNAi (1)
C
PC3
≥15 yH2AX foci [%]
Aciclovir
UL97 RNAi (1)
D
12d c-caspase-3+ cells [%]
0 µM aciclovir
300 µM aciclovir
VCaP LAPC-4 LNCaP DU145 PC3 LREX'
E
12d c-caspase-3+ cells [%]
0 µM GCV
100 µM GCV
p=0.06
VCaP LAPC-4 LNCaP DU145 PC3 LREX'
F
PC3
c-caspase-3+ cells [%]
Aciclovir
UL97 RNAi (1)
G
PC3
c-caspase-3+ cells [%]
Aciclovir
IE1/2 RNAi (1)
H
LNCaP
Ki-67+cells [%]
c-caspase-3+ cells [%]
p=0.19
vehicle
enzalutamide
enzalutamide 5d (+7d) ACV
I
LAPC-4
c-caspase-3+ cells [%]
12d
5d (+7d)
p=0.10
ns
Aciclovir (µM)
J
PC3
c-caspase-3+ cells [%]
12d
5d (+7d)
ns
Aciclovir (µM)
K
LNCaP
EdU+ cells [%]
pS10-hH3+ cells [%]
ns
p=0.14
Aciclovir
0 µM
300 µM
7d
3d (+4d)
L
300 µM Aciclovir
LNCaP
yH2AX
12d
5d (+7d)
≥15 yH2AX foci [%]
≥20 yH2AX foci [%]
12d
5d (+7d)
LNCaP PC3
M
s-aciclovir [µM]
25 mg/g food
valaciclovir
N
PC3 xenografts 17-21 days treatment
c-caspase-3+ cells [%]
control
25 mg/g valaciclovir
c-caspase 3
control
valaciclovir
O
Mouse Anterior Prostate
cleaved caspase-3
Control
Valaciclovir

**Supplemental Figure 6: Pre-clinical evaluation of aciclovir**

**A)** Aciclovir (300 μM) induced DNA damage in LNCaP, LAPC-4 and PC3, examined after 12 days by quantification of γH2AX foci per cell (n=3). Scale bar 20 μm. **B)** In PC3, UL97 RNAi (1) reduced the ability of ganciclovir (GCV) to promote γH2AX foci (n=3), examined after 7 days. **C)** In PC3, UL97 RNAi (1) reduced the ability of aciclovir (300 μM) to promote γH2AX foci (n=3), examined after 7 days. **D)** Aciclovir (300 μM) induced apoptosis (cleaved caspase-3) in cell lines examined after 12 days (n=3-4). **E)** Ganciclovir (100 μM) induced apoptosis (cleaved caspase-3) in several cell lines examined after 12 days (n=3-5). **F)** In PC3, UL97 RNAi (1) reduced the ability of aciclovir to induce apoptosis (n=3), examined after 7 days. **G)** In PC3, IE1/2 RNAi (1) reduced aciclovir induced apoptosis (n=3), examined after 7 days. **H)** Enzalutamide reduced the percentage of Ki-67$^+$ cells in LNCaP and reduced the ability of aciclovir (ACV) to induce apoptosis (cleaved caspase-3) (n=3). **I-J)** LAPC-4 and PC3 treated with aciclovir in different concentrations continuously for 12 days or 5 days with and 7 days without (5d (+7d)) (n=3). **K)** LNCaP treated with aciclovir and treated with EdU 1 hour prior to fixation. Percentage of EdU$^+$ cells and pS10-hH3$^+$ cells were quantified (n=3). **L)** Discontinuous aciclovir treatment increased γH2AX foci in LNCaP and PC3 (n=3-4). Scale bar 20 μm. **M)** After seven to nine days of valaciclovir treatment, serum aciclovir levels were analyzed. Box plot (median, 25-75$^{th}$ percentiles, error bars show min-max values). **N)** Mice were untreated (control, n=4) or treated with valaciclovir (n=3) 17-21 days. Aciclovir induced apoptosis (cleaved caspase-3$^+$ cells) in PC3 xenografts. Percentage of cleaved caspase-3$^+$ cells were compared with unpaired t-test. Scale bar: 25 μm. **O)** In prostates of mice treated with valaciclovir 17-21 days, no induction of apoptosis (cleaved caspase-3$^+$ cells) was observed. Scale bar: 100 μm. Cell nuclei are labeled in blue with DAPI. Data in bar graphs are shown as mean. Error bars in bar graphs are SEM. Data are analyzed with paired t-tests unless otherwise specified. p<0.05 is*, p<0.01 is **. p<0.001 is ***. Ns is non-significant. In (A)-(C) and (L), percentage of cells with 15 or 20 or more foci were analyzed and shown in graphs.

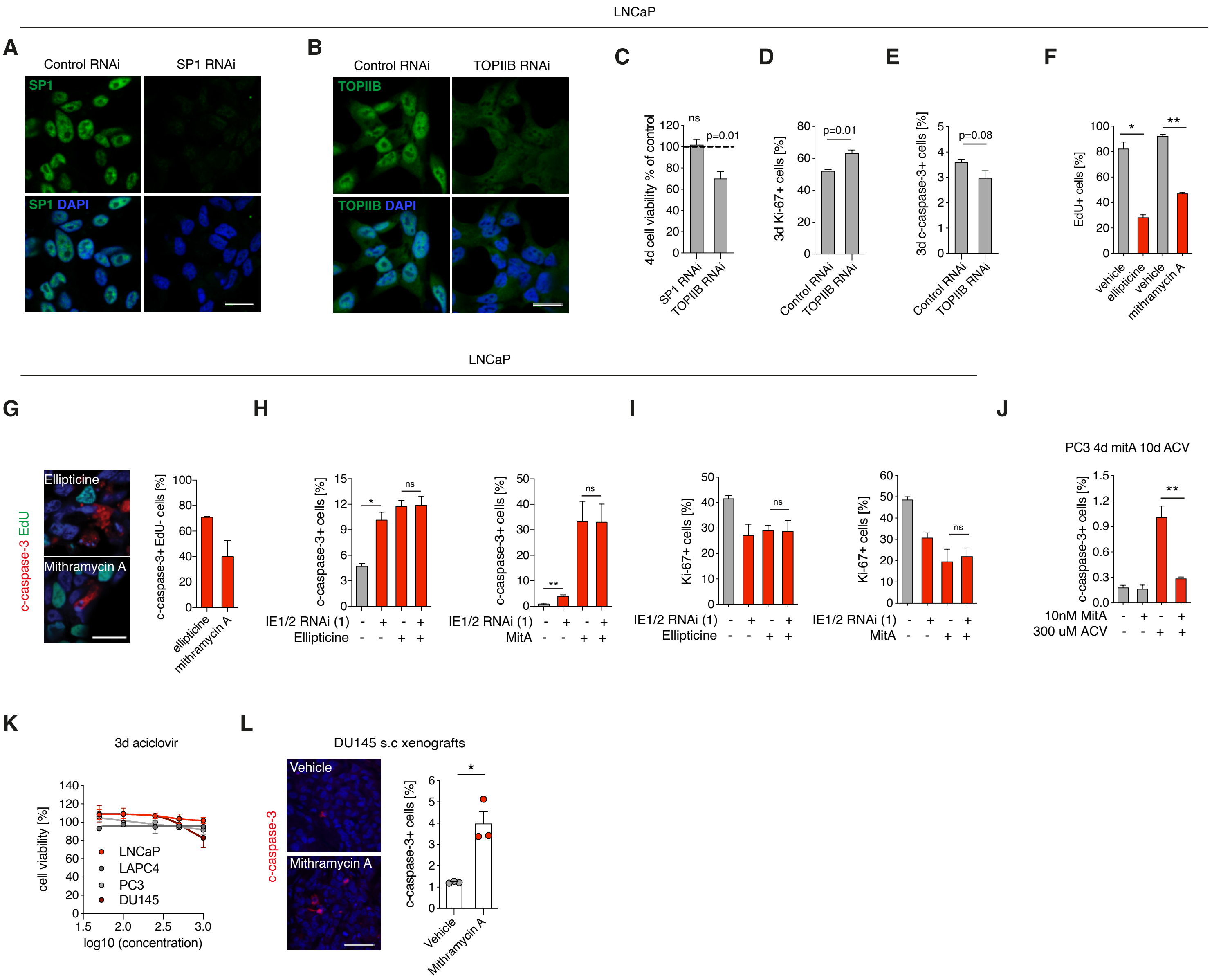
LNCaP
A
Control RNAi
SP1 RNAi
SP1
SP1 DAPI
B
Control RNAi
TOPIIB RNAi
TOPIIB
TOPIIB DAPI
C
4d cell viability % of control
ns
p=0.01
SP1 RNAi
TOPIIB RNAi
D
3d Ki-67+ cells [%]
p=0.01
Control RNAi
TOPIIB RNAi
E
3d c-caspase-3+ cells [%]
p=0.08
Control RNAi
TOPIIB RNAi
F
EdU+ cells [%]
vehicle
ellipticine
vehicle
mithramycin A
LNCaP
G
c-caspase-3 EdU
Ellipticine
Mithramycin A
c-caspase-3+ EdU- cells [%]
ellipticine
mithramycin A
H
c-caspase-3+ cells [%]
IE1/2 RNAi (1)
Ellipticine
MitA
I
Ki-67+ cells [%]
IE1/2 RNAi (1)
Ellipticine
MitA
J
PC3 4d mitA 10d ACV
c-caspase-3+ cells [%]
10nM MitA
300 uM ACV
K
3d aciclovir
cell viability [%]
LNCaP
LAPC4
PC3
DU145
log10 (concentration)
L
DU145 s.c xenografts
c-caspase-3
Vehicle
Mithramycin A
c-caspase-3+ cells [%]
Vehicle
Mithramycin A

**Supplemental Figure 7: Therapeutic targeting of CMV in prostate cancer is not dependent on SP1 and TOPIIB**

**A-B)** SP1 and TOPIIB protein was reduced upon transfection with RNAi in LNCaP. **C)** Cell viability shown as percentage of control four days after transfection with SP1 RNAi or TOPIIB RNAi in LNCaP (n=3). **D-E)** Percentage of Ki-67+ cells and cleaved caspase-3$^+$ cells were evaluated in LNCaP transfected with Control RNAi or TOPIIB RNAi after three days (n=3). **F)** Percentage of EdU$^+$ cells in LNCaP treated continuously with EdU and ellipticine (3μM) for one day or mithramycin A (10 nM) for three days (n=3). **G)** Cells were positive for the apoptosis marker cleaved caspase-3 (n=3) independent of EdU incorporation. **H-I)** LNCaP was transfected with Control RNAi or IE1/2 RNAi (1) four days in total and co-treated with ellipticine for one day and mithramycin A for three days (n=3). Percentage of cleaved caspase-3 and Ki-67$^+$ cells are shown in bar graphs. **J)** In PC3, pre-treatment with mithramycin A four days (10 nM) reduced aciclovir induced apoptosis (n=3). **K)** Cell viability in LNCaP, LAPC-4, PC3 and DU145 after three days of 0-1000 μM aciclovir (n=3), shown in log scale. **L)** Percentage of cleaved caspase-3$^+$ cells in DU145 xenografts in vehicle and mithramycin A treated animals (n=3 per treatment, all datapoints are shown in graph). Treatments were compared with un-paired student's t-test. Scale bar: 50 μm. Cell nuclei are labeled in blue with DAPI. Data in bar graphs are shown as mean. Error bars in bar graphs are SEM. Paired t-tests were performed in (C)-(F), (H)-(J). Scale bar is 25 μm in (A), (B) and (G). $p<0.05$ is*, $p<0.01$ is **. Ns is non-significant.

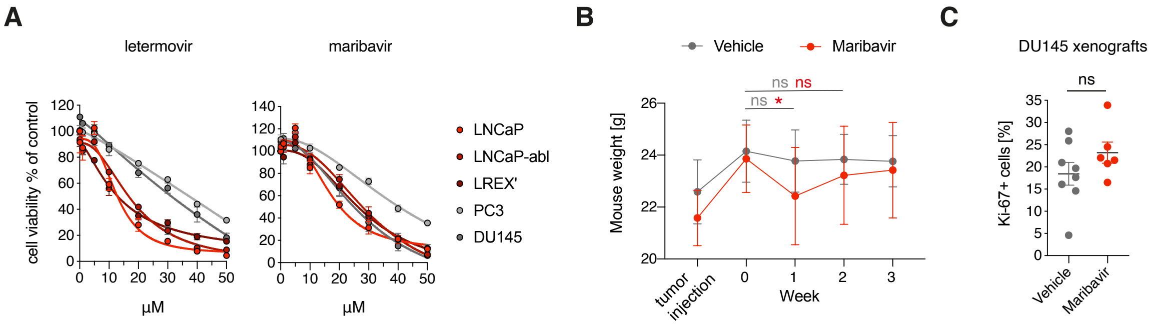
A
letermovir
maribavir
cell viability % of control
μM
LNCaP
LNCaP-abl
LREX'
PC3
DU145
B
Vehicle
Maribavir
Mouse weight [g]
ns ns
ns *
tumor injection
Week
C
DU145 xenografts
Ki-67+ cells [%]
ns
Vehicle
Maribavir

**Supplemental Figure 8: Letermovir and maribavir in prostate cancer models**

**A)** Cell viability examined after six days of treatment with letermovir (0-50 μM) or maribavir (0-50 μM) (n=3-4). Lines show non-linear fit of treatment in each cell line. **B)** Animal weight was examined at time of tumor injection and weekly at start of treatment with vehicle (n=8) or maribavir (n=10). Error bars are SD. Unpaired two-tailed student's t-tests were performed comparing 0-1 weeks and 0-2 weeks for vehicle (grey) and maribavir (red) treated mice. **C)** Ki-67+ cells in tumors of mice treated with vehicle (n=8) or maribavir (n=6) 3.5 weeks. Line shows mean. Error bars are SEM. Unpaired two-tailed student's t-test was performed. $p<0.05$ is*, ns is non-significant.

**Table S1.**
**Characteristics of aciclovir epidemiology cohort**

| | Aciclovir non-users<br>N (%) | Aciclovir users<br>N (%) |
|---|---|---|
| Overall | 780,580 (100) | 156,116 (100) |
| **Age at index date (years)** | | |
| Mean (SD) | 61.5 (12.7) | 61.5 (12.7) |
| **Calendar year** | | |
| 1996-2000 | 94,130 (12.1) | 18,826 (12.1) |
| 2001-2005 | 132,945 (17.0) | 26,589 (17.0) |
| 2006-2010 | 167,505 (21.5) | 33,501 (21.5) |
| 2011-2015 | 186,550 (23.9) | 37,310 (23.9) |
| 2016-2020 | 199,450 (25.6) | 39,890 (25.6) |
| **Charlson Comorbidity Index** | | |
| Low (comorbidity score = 0) | 550,099 (70.6) | 88,786 (63.9) |
| Medium (comorbidity score 1 or 2) | 176,268 (22.6) | 39,992 (25.6) |
| High (comorbidity score 3+) | 53,413 (6.8) | 16,338 (10.5) |

### ONLINE METHODS

#### Post-mortem donors

Ethical permission for study of human samples was granted by the regional ethics committee of Sweden (2010/313-31/3). Prostate tissue and blood samples were collected from post-mortem donors between 2015 and 2020 through KI Donatum, Karolinska Institutet, Stockholm, Sweden, as described in[1]. We collected cross sections of prostate tissue that were designed to include parts of the central-, transition- and peripheral zone of the gland from 41 male post-mortem donors aged 19-89. One side of the prostate was frozen at -80°C and one side was subjected to fixation and paraffin embedding. Serum was separated from other blood products by centrifugation at 20 minutes, 3500 rpm at 4 degrees. Histological evaluation of prostate cancer was assessed in H&E stained FFPE prostate slides by a trained pathologist. An incidental prostate cancer (Gleason score 3+3) was detected in a subject and a suspected incidental prostate cancer was detected in another subject but was difficult to further assess histologically due to post-mortal effects on the tissue. Presence of cancer was validated by lack of p63 protein expression by immunohistochemistry. Two additional individuals had been treated for prostate cancer with local radiation or chemotherapy. Medical history was obtained from medical journals, next of kin, police reports and patient registry, all in accordance with ethical permit 2010/313-31/3.

#### Material from prostate cancer patients

De-identified prostate cancer FFPE tissue from prostatectomy specimens with matched plasma was received from the Prostate Cancer Biorepository Network (PCBN) biobank. The locations of tumors were outlined in H&E stained sections by a pathologist associated with PCBN. Ethical permission for biobank tissue collection was obtained from the local institutional review board at Memorial Sloan Kettering Cancer Center, New York, USA. A human prostate tissue array (cancer) (NBP2-30169; Novusbio) was used to examine an association between CMV and androgen receptor protein expression. Fresh frozen and FFPE bone metastasis samples from patients with castration resistant prostate cancer were collected at Umeå University Hospital (cohort described in e.g.,[2]). Collection and analyses of bone metastases were approved by the local ethics review board of Umeå University (Dnr 2016-332-32M). Phenotype characterization of bone metastases, divided into MetA (high AR activity), MetB (dedifferentiated, increased proliferation) and MetC (non-luminal, EMT enriched) was described previously[3]. Analysis of CPC-GENE ultra-deep RNA-sequencing data has partly been described in[1]. In addition, reads mapping to a CMV genome (NC_006273.2), which was concatenated with the human genome, were explored (n=144). Default parameters with STAR was used to align reads. No CMV specific hits were detected when analyzing 173 CMV genes. Contaminant reads mapping to the CMV Major Immediate Early exclusive to the CMV genome were recovered in 19 of 144 samples.

**Aciclovir population cohort**

Danish law does not require approval from an ethics committee or informed consent from patients to perform registry-based studies. We conducted a cohort study based on data from the following population-based Danish registries: the Cancer Registry[4], The Cause of Death Registry, the Danish National Prescription Registry[5], the Danish National Patient Registry[6], and the Danish Civil Registration System[7]. Unambiguous linkage between Danish registries is possible using the Danish Civil Registration Numbers assigned to all Danish residents since 1968, at birth, or upon immigration to the country[7]. Danish citizens have equal tax-supported access to health care provided by the Danish National Health Service[8]. The Danish Cancer Registry has recorded incident cases of cancer on a nationwide basis since 1943 and has been shown to have an almost complete case ascertainment. Cancer diagnoses in the Cancer Registry are recorded according to the International Classification of Diseases, version 10, and the International Classification of Diseases for oncology for topography and morphology codes. Prostate cancer was defined by ICD-10 code C61 in the Cancer Registry. Prostate cancer specific mortality was recorded in the Cause of Death registry by ICD-10 code C61. We retrieved information available on prostate cancer incidence and prostate cancer specific mortality from 1970 to 2020.

The Danish National Prescription Registry contains information on all prescriptions dispensed at community pharmacies in Denmark since 1995[5]. For each prescription, the Prescription Registry records date and full description of the dispensed product including the Anatomical Therapeutic Code (ATC). We retrieved information available from the Prescription Registry from 1995 to 2020. We identified males prescribed systemic aciclovir (aciclovir ATC J05AB01 or valaciclovir ATC J05AB11) between 1996-2020 and used date of first prescription as start of follow-up (n = 313,072). Follow up starts one year after the initiation of the Danish National Prescription Registry (1996 and 1995 respectively). Due to the Danish National Prescription Registry not existing prior to 1995, it is unknown if men have been prescribed systemic aciclovir prior to this year. This can result in misclassification, and we partly alleviate this bias by having at least one year look back for each study participant to find actual new drug users and matched non-users. Systemic aciclovir indications include shingles (varicella zoster reactivation), prophylaxis against genital herpes (herpes simplex 2 reactivation) and prophylaxis against CMV reactivation in patients with transplanted organs or stem cells.

Next, we applied several exclusion criteria: males below age 40 at their first prescription were excluded; n = 144,352 (46.1%). Prevalent aciclovir users (determined from 1995 and onwards) were excluded; n = 7,628 (2.4%). Males prescribed with other nucleotide analogues before start of follow-up were excluded; n = 460 (0.1%). Males with prostate cancer before start of follow-up were excluded; n = 4,516 (1.4%). Therefore, the final cohort of aciclovir users consisted of 156,116 men. We also required that matched non-users had no prostate cancer diagnosis and no ATC J05AB (nucleotide analogues including aciclovir and valaciclovir) prescription prior to their index date (the date of first prescription for the corresponding user). Aciclovir users were matched 1:5 to systemic aciclovir/valaciclovir non-users from the general population (from here defined as aciclovir non-users). Aciclovir users were matched to non-users on year of birth and calendar year grouped by four-year intervals (Table S1). In total, the study cohort included 780,580 aciclovir non-users (Figure S10). Calendar year group was used as a matching factor to account for differences in s-PSA screening with time.

The mean age for aciclovir users was 61.5 (SD, 12.7; Table S1) and study participants were included throughout 1996 to 2020 (calendar year, grouped by 4 years; Table S1). The median follow-up time was 7.6 years (IQR, 3.5-12.8) years with 1,324,582 person-years for aciclovir non-users and 6,744,409 person-years for aciclovir users (Table S1). During follow-up, 36,428 men (3.9%) were diagnosed with prostate cancer and 8,366 men died from prostate cancer (0.9% aciclovir non-users, 0.8% aciclovir users) (Table S1). Aciclovir users were more likely to have a high Charlson Comorbidity Index (three or higher) (Table S1).

**CMV Ig serum analysis**
Anti-CMV IgG titer in serum was analyzed using CMV IgG, CMIA assay (Architect, Abbott) or by CMV IgG CLIA (chemiluminescence immunoassay) on the LIAISON®XL Analyzer at Karolinska University Hospital Clinical Microbiology Laboratory.

**Western blot human samples and cells for CMV proteins**
Frozen human prostate tissue was cut in 10-30 µm thin sections on a cryostat into tubes. Lysis buffer (20mM Tris pH 7,5, 1% triton X-100, 150mM NaCl, 5mM EDTA, 10% glycerol, 10mM NaF) with Halt protease & phosphatase inhibitor cocktail (78440, ThermoFisher) was added to samples. Prostate lysates were treated with plastic pestles. Cell lysates and prostate lysates were vortexed 30 seconds and were then centrifuged for 10 minutes at 16,9 RCF at 4ºC and supernatants were collected. Protein concentration was quantified with Pierce BCA Protein Assay Kit (23225, ThermoFisher) according to manufacturer's instructions. Prostate protein samples (40 µg) and cell protein samples (10-20 µg) were incubated in NuPAGE LDS sample buffer (NP0007, ThermoFisher) with 10% B-mercaptoethanol according to manufacturer's instructions. Samples were loaded onto NuPAGE 4 to 12%, Bis-Tris, 1.5 mm, Mini Protein Gel, 10-well (NP0335BOX, ThermoFisher) and electrophoresis was performed in NuPAGE MOPS SDS running buffer (NP0001, ThermoFisher) with NuPAGE antioxidant (NP0005, ThermoFisher). BenchMark Pre-stained Protein Ladder (10748010, ThermoFisher) or PageRuler Prestained protein ladder (26616, ThermoFisher) was used as a protein ladder. After electrophoresis, proteins were transferred to Trans-Blot Turbo Mini 0.2 µm PVDF Transfer Packs (1704156, Biorad) using a trans-blot turbo transfer system (Biorad). Membranes were incubated in Superblock blocking buffer in TBS (37535, ThermoFisher) with 0,05% tween-20 for minimum of 30 minutes for CMV protein blots. For non-CMV protein blots, membranes were blocked in milk.

Primary antibodies were diluted in blocking buffer and incubated over night at 4°C. Primary antibodies used: Cytomegalovirus US28 (1:2000, rabbit, PA5-102302, ThermoFisher), Cytomegalovirus UL97 (1:2000, rabbit, PA5-99784, ThermoFisher), Androgen Receptor (1:2000, mouse, 411, sc-7305, Santa Cruz). As an endogenous protein level control, B-actin (1:5000, mouse, clone AC-74, A2228, SigmaAldrich) was used and incubated for one hour. Membranes were washed in TBS with 1% tween-20 (TBS-T) and incubated with secondary antibodies diluted in blocking buffer for two hours (1:5000, anti-mouse-HRP, NA931V; anti-rabbit-HRP, NA934V, GE healthcare). After washing in TBS-T, membranes were developed with SuperSignal west dura extended duration substrate (ThermoFisher) or SuperSignal West Atto (ThermoFisher) and imaged on a Chemidoc MP imaging system (Biorad). Non-CMV protein blots were developed with ECL and films.

Immunoblot bands were quantified in ImageJ64 with a gel analysis tool using B-actin as loading control.

**CMV IHC on human FFPE tissues**
Tissues were fixed in formaldehyde, embedded in paraffin (FFPE) and cut to slides at a thickness of 4 μm. FFPE tissue slides were deparaffinised and rehydrated in xylene and an ethanol gradient. After a wash in dH2O, slides were incubated in 1X Antigen Unmasking Solution, Citric Acid Based (100X, H-3300, Vector Laboratories) 30 minutes in a steamer, cooled for 10 minutes and then washed in PBS. CMV positive control slides (CSC0925P, American MasterTech) were incubated in antigen unmasking solution for 20 minutes in a steamer. Slides were incubated for 10-15 minutes in BLOXALL blocking solution (SP-6000, Vector Laboratories) or 3% $H_2O_2$, washed in PBS, incubated 20 minutes with FC receptor blocker (Innovex biosciences), incubated 45 minutes with 10% donkey serum containing 0,5% triton, and then incubated with avidin/biotin blocking kit (Vector Laboratories) according to manufacturer's instructions.

Primary antibodies were then diluted in 10% donkey serum and incubated at room temperature overnight. Primary antibodies used: Cytomegalovirus US28 (rabbit, 1:150, PA5-39864, polyclonal, ThermoFisher), Cytomegalovirus pp65 (mouse, 1:50, clone CH12, sc-56976, monoclonal, Santa Cruz Biotechnology), Cytomegalovirus pp71 (goat, 1:200, clone vC-20, sc-33323, polyclonal, Santa Cruz Biotechnology), Cytomegalovirus IE1 (mouse, 1:10, pp72, clone 6E1, sc-69834, Santa Cruz Biotechnology), Cytomegalovirus IE1/2 (mouse, 1:150, MAB810R, monoclonal, Millipore), Chromogranin A (mouse, 1:100, LK2H10, MA5-13096, ThermoFisher), keratin 5 (rabbit, 1:200, EP1601Y, ab52635, Abcam), keratin 18 (rabbit, 1:100, H-80, sc-28264, Santa Cruz), TP63 (mouse, 1:100, clone 4A4, CM163A, Biocare Medical), Ki-67 (rabbit, 1:100-250, clone SP6, ThermoFisher), wide spectrum cytokeratin (rabbit, 1:100, ab9377, abcam), pan-keratin (mouse, 1:200, clone C11, #4545, Cell Signaling), AR (rabbit, 1:50-100, clone SP107, ThermoFisher).

After washing in PBS, slides were incubated with a secondary antibody conjugated with biotin (donkey anti mouse-biotin or donkey anti rabbit-biotin, 1:500, Jackson laboratories) or a secondary antibody conjugated with a fluorophore (donkey anti mouse 488/cy3/cy5, donkey anti rabbit 488/cy3/cy5, Jackson laboratories). If incubated with a secondary antibody conjugated with biotin, slides were then incubated with VECTASTAIN Elite ABC HRP Reagent, R.T.U. (PK-7100, Vector laboratories) 30 minutes, after which slides were washed in PBS and incubated with ImmPACT DAB (Vector laboratories) or TSA using Alexa Fluor 488 tyramide SuperBoost kit (B40932, ThermoFisher) according to manufacturer's instructions. For co-labelling, slides were treated with 3% hydrogen peroxide for 15 minutes, incubated with primary antibody overnight, and was then developed with TSA Tyramide 555 or Tyramide 647. DAB-stained slides were counterstained with hematoxylin QS (Vector laboratories) after washing in water, dehydrated in an ethanol series from 50% to 100%, incubated in xylene, air dried and mounted with pertex mounting medium (Histolab). Fluorescently stained slides were counterstained with DAPI (1:5000, SigmaAldrich) and mounted with Prolong Gold Antifade mounting medium (ThermoFisher).

**CMV DNA FISH on human prostate tissue**

FFPE prostate tissue slides were deparaffinized and rehydrated in xylene and ethanol gradient. After a wash in dH2O, slides were incubated in 1X RISH Retrieval (RI0209M, Biocare Medical) 15 minutes and were thereafter cooled for 10 minutes. Slides were incubated with 3% H2O2 in methanol 5 minutes and washed in dH2O, incubated in 1:4 RISHzyme in RISHzyme buffer for 1 minute, washed in dH2O and then incubated in DNAse free RNAse (ROSCH) 1:5 in 2x SSC buffer 30 minutes at 37$^{o}$C. Slides were incubated with the digoxigenin labelled RISH CMV probe (RI0011T) 15 minutes 95$^{o}$C after which slides were incubated with probe overnight at 37$^{o}$C. Hybridised slides were then subjected to stringency washes. Slides were washed in 4x SSC 5 minutes x2 at room temperature and 0,01x SSC 5 minutes x2 80$^{o}$C. Slides were incubated with secondary reagent and tertiary reagent per protocol (RISH HRP detection kit, Biocare Medical) after which CMV hybridization was developed with TSA using Alexa Fluor 488 tyramide SuperBoost kit (B40932, ThermoFisher) according to manufacturer's instructions. For co-labelling, slides were treated with 3% hydrogen peroxide for 15 minutes. After washing in PBS, slides were stained as described above for IHC. IHC staining was developed with TSA using Tyramide 555 or Tyramide 647.

**PBMC CD14$^{+}$ enrichment and validation**

Peripheral blood from post-mortem donors was mixed with PBS-EDTA (2 mM EDTA) and centrifuged at 900g 20 minutes in a blood separation tube containing 16 ml Lymphoprep$^{TM}$ (STEMCELL Technologies). White blood cells were separated out, washed with PBS-EDTA and run through a 100 µm filter. After 1500 g 5 minutes centrifugation, the pellet was resuspended in 1 ml MACS buffer (PBS-EDTA, 0,5% (w/w) BSA) and incubated in 1:20 CD14 MicroBeads (Miltenyi Biotec) 20 minutes at 4$^{o}$C. The suspension was washed in MACS buffer, centrifuged and resuspended in 1 ml MACS buffer. CD14⁺ cells were enriched using LS columns (Miltenyi Biotec) on a MACS® manual separator (Miltenyi Biotec). To validate the bead-enrichment, cells were incubated with phycoerythrin (PE)-conjugated anti-CD11b monoclonal antibody (1:20, BioLegend, clone ICRF44) 30 minutes at room temperature, centrifuged, and resuspended in MACS buffer. Bead-enriched cells and non-bead-enriched cells (flow-through) were analysed by flow cytometry on an INFLUX machine. FlowJo version 10.5.3 was used to analyse FACS plots and provide statistics on $CD11b^+$ cell fractions, after gating out cell debris. Bead-enriched cells were stored in PBS at -80°C until further analysis.

**DNA extraction and quantitative CMV PCR**

Frozen tissue was cut into DNAse free eppendorf tubes in a clean cryostat in which the knife was changed and cryostat cleaned with ethanol between each specimen, in order to reduce the risk of contamination. Tissues and cell pellets were stored at -80°C until DNA extraction. DNA was extracted using Dneasy blood and tissue kit (QIAGEN) using manufacturer's instructions including treatment of samples with RNAse A (QIAGEN). DNA concentration was measured on a nanodrop or on Qubit using Qubit dsDNA BR Assay Kit (ThermoFisher) according to manufacturer's instructions. Amplirun cytomegalovirus DNA control (MBCO16, Vircell) was used as a positive control.

Positive control DNA was reconstituted in 100 µl buffer as per manufacturer's instructions and diluted 1:10 in dH2O. Of this, 1 µl was used per qPCR reaction using taqman fast master mix (ThermoFisher) and taqman primer/probes described below. The restriction enzyme *BsrI*, has 387 restriction sites in the CMV strain Merlin. 1000 ng prostate DNA or 300-500 ng bone metastasis DNA was incubated in a 25 µl reaction containing 1X NEBuffer 3.1 (NEB) with or without 0,2 µl of the restriction enzyme Bsr1 (10,000 units/ml; NEB) at 65deg for 16h and then 85deg 20min. For samples treated with bsr1, 5 µl was loaded into one qPCR reaction. Otherwise, input amount of DNA was 500 ng per reaction for cell lines and post-mortem prostate DNA samples and 100-200ng for $CD14^+$ cells. For cell lines in experiments, 10ng DNA was used. qPCR was performed in 96-well plates and run on a 7500 Fast Real-Time PCR system (Applied Biosystems) or CFX96 system (Biorad) with 60-70 cycles. The following master mixes were used: taqman fast master mix (ThermoFisher); UL83, GoTaq Probe qPCR master mix (Promega); UL83, UL37, and qRT-PCR Brilliant III Probe Master Mix with ROX (Agilent); UL32, UL36, UL38, UL87, UL97, UL122). The pre-designed taqman primer/probe GAPDH (Hs02786624_g1) was used.

Several custom taqman primer/probes were used:

| | |
|---|---|
| UL32 | F: GGCGCGGGAACCTCTT |
| | R: CCGTGGGCGACAAAACG |
| | Probe: CAGCCGTCAGCCTCG |
| UL36 | F: GAAAGAAGGGACACCGAAACCA |
| | R: GACAGGTGGGTGTCTTTTCCA |

| | |
|---|---|
| | Probe: ACGCACGATGGCCTC |
| UL37 (X1) | F: GTGGCCGCGCTCTTG |
| | R: GCTCTGTGTCCTCCGTTACG |
| | Probe: CCTCCCCGGCCTCG |
| UL38 | F: CGCTCCCACGTCCGT |
| | R: AGCTGGTGGAAGACCATCAC |
| | Probe: CAGCACGCGCACACTA |
| UL83 | F: CCCAGCGTGACGTGCATAA |
| | R: AGGTGTACCTGGAGTCCTTCTG |
| | Probe: CTCCGGCAAGCTCT |
| UL87 | F: GTGCTGTTTCTGCGTGCTT |
| | R: CAACTGCAGCCGCTTCTC |
| | Probe: TCGACCGTGCAGCTTG |
| UL97 | F: ACCGTCTGCGCGAATGTTA |
| | R: GTCGCAGATGAGCAGCTTCT |
| | Probe: CCACCCTGCTTTCCG |
| UL122 | F: GCTTGATGTCTTCCTGTTTGATGAG |
| | R: ACGCGTCCTTTCAAGGTGATTATTA |
| | Probe: CCTCCCGCGCCTATC |

Size of PCR products were visualized on a 3% agarose gel with 50bp ladder. The correct identity of PCR products in prostate tissues was validated with sanger sequencing on PCR products cloned into a plasmid with the TOPO TA Cloning Kit (K457501, ThermoFisher Scientific).

**Cell line culture conditions**

LNCaP and PC3 were purchased from ATCC and cultivated in RPMI-1600 (Thermo Fisher) with 10% FBS and 1% Penicillin-Streptomycin (PS). MyC-CaP was purchased from ATCC and cultivated in DMEM (ThermoFisher) with 10% FBS and 1% PS. LAPC4 was a gift from Robert Reiter, University of California, Los Angeles, USA. LAPC4 was cultivated in IMDM with 5% FBS and 1% PS. LREX' (Arora et al., 2013) was a gift from Charles Sawyers, MSKCC, New York, USA. LREX' was cultivated in 20% FBS, 1 µM enzalutamide and 1% PS. LNCaP-abl (Culig et al., 1999) was a gift from Helmut Klocker, Medical University of Innnsbruck, Austria. LNCaP-abl was cultivated in RPMI-1600 with 10% charcoal stripped FBS and 1% PS. Since their arrival to our laboratory, the cell lines have been handled and cultured in the absence of purified CMV, minimizing risk of contamination from laboratory strains.

**CMV DNA FISH on xenografts and mouse prostate**

Deeply sedated animals were perfused with 4% formaldehyde and were post-fixed over night. The prostate glands and DU145 xenografts were dissected and incubated in sucrose 30% w/w at 4$^{o}$C. Tissue was placed in OCT and cut to glass slides on a cryostat at 12 $\mu$m thickness. Tissue slides were stored at -20$^{o}$C. Slides were warmed at 42$^{o}$C for 1 hour. Slides were treated with 3% $H_2O_2$ diluted from 30% in methanol for

5 minutes. Slides were washed in $dH_2O$ and treated with RISHzyme (1:4 mouse prostate OCT; 1:8 LNCaP xenografts OCT) in RISHzyme buffer (Biocare Medical) for 1 minute. Slides were then incubated with RNAse, DNAse free (Roche, Sigma Aldrich) 1:5 in 2x SSC buffer at 37$^oC$ for 30 minutes. RNAse was washed off and slides were incubated with a digoxigenin labelled CMV probe (RISH, Biocare Medical) at 95$^oC$ for 5 minutes and then at 37$^oC$ overnight. Hybridized slides were then subjected to stringency washes. Slides were washed in 4x SSC 5 minutes x2 at room temperature and 0,01x SSC 5 minutes x2 80$^oC$. Slides were incubated with secondary reagent and tertiary reagent per protocol (RISH HRP detection kit, Biocare Medical) after which CMV hybridization was developed with TSA coupled to Cy3 (NEL744001KT, PerkinElmer) for 10 minutes. Slides were then incubated with 10% NDS containing 0,5% triton 45 minutes after which slides were incubated over night with an antibody against wide spectrum cytokeratin (1:100, rabbit, ab9377, abcam) diluted in 10% NDS. After washing in PBS, slides were incubated with a secondary antibody 1 hour (donkey-anti-rabbit-488, 1:500, Jackson laboratories), after which slides were stained with DAPI (1:5000, sigma) and mounted with prolong gold mounting medium (ProLong™ Gold Antifade Mountant, ThermoFisher).

**Prostate cancer cell transfection**

Cells were transfected with plasmids using Lipofectamine 3000 transfection reagent according to manufacturer's instructions (Cat. No. L3000001, ThermoFisher). HCMV UL97 plasmid, tagged with HA [9], was a gift from Robert Kalejta (Addgene plasmid # 26687). pcDNA 3.1 (+) mammalian expression vector (ThermoFisher) was used as control plasmid.

Cells were treated with RNA interference with Lipofectamine RNAiMax transfection reagent (Cat. No. 13778030, ThermoFisher) according to manufacturer's instructions. For stealth siRNA, stock solutions of siRNA used at 20 µM and for silencer select siRNA stock solutions of siRNA at 10 µM were used. As control, scrambled siRNA was used; Stealth RNAi siRNA Negative Control, Med GC (ThermoFisher Scientific) or Silencer Select Negative Control #1 siRNA (ThermoFisher Scientific). Stealth siRNA against SP1 and TOPIIB was used. Custom designed Stealth siRNA and Silencer Select siRNA (ThermoFisher Scientific) were used to target CMV gene expression. Silencer select siRNA were used against CMV-IE1/2 (2) and CMV-IE1 (2) with sequences used from[10].

| | |
|---|---|
| SP1 | 5' GCAGACACAGCAGCAACAAAUUCUU '3 |
| | 5' AAGAAUUUGUUGCUGCUGUGUCUGC '3 |
| TOPIIB | 5' CCAGCAUGAUGAUAGUUCCUCCGAU '3 |
| | 5' AUCGGAGGAACUAUCAUCAUGCUGG '3 |
| CMV-IE1/2 (1) | 5' ACCUUUGAACAAGUGACCGAGGAUU '3 |
| | 5' AAUCCUCGGUCACUUGUUCAAAGGU '3 |
| CMV-IE1/2 (2) | 5' GGAAGGAGGUUAACAGUCAUU '3 |
| | 5' UGACUGUUAACCUCCUUCCUU '3 |

| | |
|---|---|
| CMV-IE1 (1) | 5' GCGGGAGAUGUGGAUGGCUUGUAUU '3 |
| | 5' AAUACAAGCCAUCCACAUCUCCCGC '3 |
| CMV-IE1 (2) | 5' GGAAGAAAGUGAACAGAGUUU '3 |
| | 5' ACUCUGUUCACUUUCUUCCUU '3 |
| CMV-UL97 (1) | 5' UCAGCGAGCCCUAUCCGGAUUACAA '3 |
| | 5' UUGUAAUCCGGAUAGGGCUCGCUGA '3 |
| CMV-UL97 (2) | 5' AAAGCAGGGUGGUAACAUUCGCGCA '3 |
| | 5' UGCGCGAAUGUUACCACCCUGCUUU '3 |

**Prostate cancer cell drug treatments**

Following drugs were administered to cell lines *in vitro*: 5-α DHT (5α-Androstan-17β-ol-3-one; A8380, Sigma Aldrich), Aciclovir Hospira, concentrate for intravenous infusions (25 mg/ml aciclovir (111 mM), sodium hydroxide 4,6 mg/ml, pH 11.3-11.5, Hospira, Pfizer), ARCC 4 (7254, Tocris), Cisplatin (2251; Tocris), Ellipticine (sc200878; Chemcruz), Etoposide (E1383; Sigma Aldrich), Enzalutamide (Santa Cruz), Ganciclovir (SigmaAldrich), Letermovir (Cayman chemical), Maribavir (MedChemTronica) and Mithramycin A (sc-200909; Santa Cruz Biotechnology). EdU (ThermoFisher Scientific) was administered the whole treatment period or 1 hour prior to cell fixation at a concentration of 10 µM.

**Analysis of in vitro experiments**

Cells were seeded in replicate or triplicate. Cell viability was measured by CellTiter-Glo 2.0 Cell Viability Assay (G9242, Promega) according to manufacturer's instructions on a luminescence reader. Relative IC50 was fitted with non-linear fit and calculated in GraphPad Prism 8. Similarly, absolute IC50 was calculated, with 0% as baseline and 100% as top restraints. For analysis of apoptosis, Caspase-Glo 3/7 Assay System (G8090, Promega) was used. Apoptosis induction was determined by dividing apoptosis luminescence with cell viability luminescence. The resulting values were then compared between treated and control cells to determine fold change in apoptosis.

**RNA extraction and RT-qPCR**

For examination of CMV gene expression in post-mortem prostates and cell lines, RNA was extracted with RNeasy Plus Mini kit (74034, QIAGEN). Prior to cDNA synthesis, RNA was treated with ezDNAse enzyme (ThermoFisher Scientific) according to manufactuer's instructions. cDNA synthesis was performed with SuperScript IV First-Strand Synthesis System (ThermoFisher Scientific) using 5 μg RNA input with either oligoDT or random hexamers at 50 or 65 $^{o}$C. Of the cDNA library, 2 μl was used as input for each RT-qPCR reaction. RT-qPCR was performed using Premix Ex Taq (Probe qPCR) (RR390L; TAKARA) according to manufacturer's instructions on a CFX96 Real Time System (Biorad) qPCR machine. Taqman primer probe GAPDH (Hs02786624_g1; ThermoFisher Scientific) was used as endogenous positive control. These custom taqman primer/probes were used:

| | |
|---|---|
| LUNA | F: CCTCGGTGGGTGGTAATCC |

| | |
|---|---|
| | R: GCGCCGTCTCCGAGTTT |
| | Probe: CTCCCGCAGTCCCC |
| UL97 | F: ACCGTCTGCGCGAATGTTA |
| | R: GTCGCAGATGAGCAGCTTCT |
| | Probe: CCACCCTGCTTTCCG |
| IEcDNA ex 1-2 | Forward: TGACGAGGGCCCTTCCT |
| | Reverse: CCTTGGTCACGGGTGTCT |
| | Probe: AAGGTGCCACGGCCCG |

For cell experiments, RNA was extracted with Rneasy Plus Mini kit (74034, QIAGEN) and cDNA was made from RNA with SuperScript VILO cDNA synthesis kit (11754050, ThermoFisher) according to manufacturer's instructions. RT-qPCR was performed using TaqMan Fast Advanced Master Mix (4444556, ThermoFisher) and taqman primer/probes. RT-qPCR was performed on a 7500 Fast Real Time PCR System (ThermoFisher) or a CFX96 Real Time System (Biorad). These taqman primer/probes were used: AR (Hs00171172_m1), KLK3 (Hs02576345_m1), TMPRSS2 (Hs01122322_m1), GAPDH (Hs02786624_g1).
AR-V7 taqman primer/probes were custom made:

| | |
|---|---|
| AR-V7 | Forward: TGTCGTCTTCGGAAATGTTATGA |
| | Reverse: TCATTTTGAGATGCTTGCAATTG |
| | Probe: TCTGGGAGAAAAATT |

**Immunofluorescence on cells and mouse tissue**
Cells grown in 8-well culture slides (BD Bioscience) were fixed in 4% formaldehyde 10-15 minutes and washed in PBS. Mice were perfused with 4% formaldehyde overnight and then washed in PBS. Xenografts and tissues were incubated in 30% sucrose and then embedded in OCT and cut at 12 μm thin sections to glass slides in a cryostat. Cells were incubated with 3% BSA with 0,3% triton or 10% Donkey serum with 0,5% triton for 20 minutes at room temperature. Tissues and xenografts were incubated with 10% Donkey serum with 0,5% triton 45 minutes. Primary antibodies were diluted in wither 1% BSA or 10% donkey serum and incubated at room temperature 3 hours or overnight at 4$^{o}$C.

Primary antibodies used: γ-H2AX (pS139) (rabbit, 1:500, ab2893, Abcam), Cleaved caspase-3 (rabbit, 1:250, clone D175, 9661S, Cell Signaling), HA-tag (rabbit, 1:250, clone C29F4, 3724P, Cell Signaling), Histone H3 (phospho S10) (rabbit, 1:500-1000, ab5176, Abcam), Ki-67 (rabbit, 1:250, clone SP6, ThermoFisher), Ki-67 (rat, 1:500-1000, clone SolA15, 14-5698-82, eBioscience), Cytomegalovirus IE1/2 (mouse, 1:200, MAB810R, Millipore), Cytomegalovirus IE1 (mouse, 1:100, pp72, clone 6E1, sc-69834, Santa Cruz Biotechnology), Cytomegalovirus UL97 (rabbit, 1:100, gift from the Coen laboratory, Harvard University), SP1 (rabbit, 1:100, 5931S Cell Signaling), TOPIIB (rabbit, 1:100, ab72334, Abcam), pRB (S807/811) (rabbit, 1:250, D20B12, 8516, Cell Signaling).

Slides were then washed with PBS and incubated with secondary antibody (Donkey anti rabbit cy3, 1:1000, Jackson Laboratories), diluted in 1 % BSA for 45-60 minutes. For EdU detection, Click-it EdU Alexa Fluor 647 Imaging Kit or Click-it Edu Plus Alexa Fluor 647 Imaging Kit (ThermoFisher Scientific) was used prior to antibody staining. Slides were washed with PBS and incubated with DAPI (1:5000, Sigma), washed in PBS and mounted with ProLong Gold Antifade Mountant (ThermoFisher Scientific).

**Microscopy and image analyses**

CMV IHC and FISH on human prostate tissue visualized with DAB was analyzed using a light microscope (CTR6000, Leica) and pictures were taken using LAS X software (Leica) with a 40x or 20x objective. CMV IHC and FISH visualized with fluorescence was analyzed using a Zeiss Imager. Z2 or a Zeiss LSM 700 confocal microscope. Pictures were taken using ZEN 2012 SP1 (8.1) software (Zeiss) and images for publication were processed in ImageJ64 or ImageJ v.1.52 and photoshop. When analyzed in fluorescence microscope, pan-keratin was used as a marker to define epithelial cells from other cell types.

For quantifications of CMV IHC abundance, a 20x objective was used to analyze a whole section of prostate tissue and 10x objective was used in analyses of bone metastases. When an area with epithelium were in sight, this was determined to be positive or negative for CMV containing epithelial cells. A CMV positive epithelial area could be completely positive for CMV or contain a lower number of CMV positive cells. When analyzed in light microscope, no marker for epithelial cells was used, as the epithelium was clearly visible with hematoxylin. For CMV-US28, varying numbers of cells with very high US28 cytoplasmic staining was found throughout tissue sections, independent of CMV serostatus. These were considered as background-stained cells and were not considered when determining CMV abundance in epithelial areas.

Intensity in cell nuclei (CMV DNA, AR protein) were compared in confocal images with ImageJ64 or ImageJ v.1.52 using the IntDen function. For quantification of percentage of cells positive for a marker, the number of cells was either counted manually or by using a cell quantification feature in ImageJ using threshold to quantify number of cells. For quantification of number of γH2AX foci per cell, 27-50 cells were analyzed per well in ImageJ64 in images taken with 40x or 63x objective.

**Animal experiments**

Animal experiments were approved by the Swedish board of agriculture, Stockholm (application 6727-18, application N132/13 with amendments N150/16 and N170/16). Adult male SCID mice (Fox Chase SCID® Mice CB17/Icr-Prkdcscid/IcrIcoCrl, Charles River Laboratories) were used.

To determine dosage of aciclovir and plasma concentrations, animals were administered with valaciclovir in food, after which blood was taken and aciclovir concentrations were measured. Valaciclovir (VALTREX, GlaxoSmithKlein) 500 mg caplets were grinded to a fine powder using pestle and mortar. Valaciclovir powder was mixed into 60-80 grams of porridge food sucrose. Concentrations of valaciclovir were 0 mg valaciclovir/g food, 25 mg valaciclovir /g food and 50 mg valaciclovir/g food. After seven - nine days, blood was drawn cardially peri-mortem during deep chloral hydrate or pentobarbital sedation. Blood was let to coagulate and was then centrifuged 10 minutes at 1500g. Serum was transferred to a 1,5 ml polypropylene tube and stored at -20$^{o}$C until analysis. Serum aciclovir concentration was measured using LC-MS/MS (routinely used to measure aciclovir levels in clinical serum samples), performed at the Clinical Pharmacology Laboratory, Karolinska University Hospital, Huddinge, Sweden.

Male SCID mice were implanted with 1,5 x $10^6$ PC3 cells or 2,0 x $10^6$ DU145 cells subcutaneously on the lower dorsal flank in 1:1 RPMI medium:matrigel (Corning Matrigel Matrix, phenol red free, Corning), in a total volume of 100 µl. Tumors were measured manually with calipers using the formula $(x*y^2)/2$. X and y are two width measurements with x being the largest measured value. When tumors had developed to mean 150 mm$^3$ in volume, animals were administered with drugs, after which animals were weighed and monitored for tumor growth weekly. Mithramycin A (750 µg/kg) was administered i.p once a day week 1. Week 2-3, mithramycin A was administered four times with at least one day intermission in between dosages. Week 4, mithramycin A was administered daily. Week 5-6, the same schedule as week 2-3 was followed. Week 7, mithramycin A was administered daily. Valaciclovir (25 mg/g food) was given in porridge food in intervals of five days with seven days intermission. Endpoint of experiment was tumor size of 1000 mm$^3$ or when humane endpoint was reached. Maribavir (100mg/kg) was given via oral gavage twice per day every weekday.

In the aciclovir xenograft experiment, deep, intradermal tumors and a tumor that was much larger at endpoint than anticipated by manual tumor measurements, were excluded from the analyses.

**Statistical analyses**

All statistical analyses were performed in GraphPad prism 8.0. All plots were also made in GraphPad prism 8.0. Two-sided unpaired t-tests were performed on numerical continuous data with normal distribution. Groups with non-normal numerical continuous data was analyzed with Mann-Whitney test. Correlation analyses on non-normal distributed data were performed with Spearman correlation analyses and correlation analyses on normal distributed data were performed with Pearson correlation analyses and with linear regression when suited. Paired/Matched values were analyzed with two-sided paired t-test. Values that were compared to a fixed value (e.g., 100% cell viability in controls) was statistically evaluated for statistical deviation from this value with one sample t-test. Proportions between two groups was analyzed with Fisher's exact test. Non-normal distributed data with more than two groups was compared with Kruskal-Wallis multiple comparisons. Survival analysis with log-rank test for animal survival in xenograft experiments was performed with Log rank test

(Mantel-Cox). Tumor size over time in mouse experiments were compared between treatment groups with repeated measures mixed-effect models, that allows for missing values, with multiple comparisons for each timepoint. Statistical analyses of the aciclovir population cohort were performed using SAS 9.4 software (SAS Institute Inc., Cary, USA). In a cox proportional hazard model, aciclovir usage, age at index date, calendar year and Charlson Comorbidity index were included as variables.

### Supplemental References

1. Classon, J.*, et al.* Prostate cancer disease recurrence after radical prostatectomy is associated with HLA type and local cytomegalovirus immunity. *Mol Oncol* (2022).
2. Ylitalo, E.B.*, et al.* Subgroups of Castration-resistant Prostate Cancer Bone Metastases Defined Through an Inverse Relationship Between Androgen Receptor Activity and Immune Response. *Eur Urol* **71**, 776-787 (2017).
3. Thysell, E.*, et al.* Gene expression profiles define molecular subtypes of prostate cancer bone metastases with different outcomes and morphology traceable back to the primary tumor. *Mol Oncol* **13**, 1763-1777 (2019).
4. Gjerstorff, M.L. The Danish Cancer Registry. *Scandinavian Journal of Public Health* **39**, 42-45 (2011).
5. Pottegard, A.*, et al.* Data Resource Profile: The Danish National Prescription Registry. *Int J Epidemiol* **46**, 798-798f (2017).
6. Schmidt, M.*, et al.* The Danish National Patient Registry: a review of content, data quality, and research potential. *Clinical Epidemiology*, 449 (2015).
7. Schmidt, M., Pedersen, L. & Sørensen, H. The Danish Civil Registration System as a tool in epidemiology. *Eur. J. Epidemiol.* **29**, 541-549 (2014).
8. Schmidt, M.*, et al.* The Danish health care system and epidemiological research: from health care contacts to database records. *Clin Epidemiol* **11**, 563-591 (2019).
9. Kuny, C.V., Chinchilla, K., Culbertson, M.R. & Kalejta, R.F. Cyclin-dependent kinase-like function is shared by the beta- and gamma- subset of the conserved herpesvirus protein kinases. *PLoS Pathog* **6**, e1001092 (2010).
10. Soroceanu, L.*, et al.* Cytomegalovirus Immediate-Early Proteins Promote Stemness Properties in Glioblastoma. *Cancer Res* **75**, 3065-3076 (2015).
